# Tracing production instability in a clonally-derived CHO cell line using single cell transcriptomics

**DOI:** 10.1101/2020.11.04.368480

**Authors:** Ioanna Tzani, Nicolas Herrmann, Sara Carillo, Cathy A. Spargo, Ryan Hagan, Niall Barron, Jonathan Bones, W. Shannon Dilmore, Colin Clarke

## Abstract

A variety of mechanisms including transcriptional silencing, gene copy loss and increased susceptibility to cellular stress have been associated with a sudden or gradual loss of monoclonal antibody (mAb) production in Chinese hamster ovary (CHO) cell lines. In this study, we utilised single cell RNA-seq (scRNA-seq) to study a clonally-derived CHO cell line that underwent production instability leading to a dramatic reduction of the levels of mAb produced. From the scRNA-seq data we identified sub clusters associated with variations in the mAb transgenes and observed that heavy chain gene expression was significantly lower than that of the light chain across the population. Using trajectory inference, the evolution of the cell line was reconstructed and was found to correlate with a reduction in heavy and light chain gene expression. Genes encoding for proteins involved in the response to oxidative stress and apoptosis were found to increase in expression as cells progressed along the trajectory. Future studies of CHO cell lines using this technology have the potential to dramatically enhance our understanding of the characteristics underpinning efficient manufacturing performance as well as product quality.

**Highlights:**

- A clonally-derived CHO cell line in our laboratory had undergone production instability – in that the amount of intact monoclonal antibody had reduced dramatically to levels at which reliable quantitation was no longer possible. We were, however, able to detect mAb heavy and light chain protein, as well as dimerised light chain species in the cell culture media.
- Single cell RNA-seq was utilised to capture > 3,800 gene expression profiles from the cell line at 72hrs post seeding.
- Analyses of the scRNA-seq data uncovered transcriptional heterogeneity and revealed the presence of multiple intra cell line clusters. The heavy chain transcript was detected at a significantly lower level in comparison to light chain transcripts. Light chain gene expression was not only more abundant, but also expressed more uniformly across the cell population.
- Using unsupervised trajectory analysis, the emergence of heterogeneity in the cell population was traced from those cells most similar to the original isolated clone to those where transcription of the mAb heavy and light chain was undetectable.
- Subsequent analysis of CHO cell gene expression patterns revealed a correlation between the progression of cells along the trajectory and the upregulation of genes involved in the cellular response to oxidative stress.

## Introduction

Recombinant therapeutic proteins such as monoclonal antibodies (mAbs) have improved the quality of life of millions of people around the world (Walsh, 2018). Tremendous progress has been made in manufacture of these medicines using Chinese hamster ovary (CHO) cells over the last 30 years and the field continues to explore new routes to improve process efficiency further (Gronemeyer, Ditz, & Strube, 2014). Recent advances in sequencing technology have enabled a considerable improvement of our understanding of CHO cell biology (Kuo et al., 2018). Next generation sequencing (NGS) has enabled the acquisition of multiple high-quality publicly available reference genomes (Kelly et al., 2017; Lewis et al., 2013; Rupp et al., 2018; Xu et al., 2011), which has in turn, facilitated the analysis of new features of CHO cell biology including DNA methylation (Wippermann, Rupp, Brinkrolf, Hoffrogge, & Noll, 2017) and epigenetics (Feichtinger et al., 2016) as well as improving mass spectrometry-based proteomics (Meleady et al., 2012). RNA-seq has also been utilised to characterise changes not only in gene expression (Clarke et al., 2019; Sha, Bhatia, & Yoon, 2018) but also alternative splicing of CHO cell mRNAs (Tzani et al., 2020). The technology has also played a crucial role in the annotation of non-coding RNA molecules such as microRNAs (Hackl et al., 2012) and long non-coding RNAs (Hernandez et al., 2019; Motheramgari et al., 2020; Vito et al., 2020). As our knowledge of the CHO cell biological system becomes increasingly sophisticated, precise routes for genetic engineering are being identified and accurate genome scale models for the prediction of cell line characteristics are being developed (Gutierrez et al., 2020).

To date, the majority of our understanding of the CHO cell transcriptome has been elucidated from the study of millions of pooled cells analysed as an individual “bulk” sample. A critical drawback of this approach is that heterogeneity, a universal characteristic of all biological systems, is ignored. Traditional bulk sample analysis provides only a “population average” limiting our understanding of complex systems, obscuring variability and in some cases describing an inferred cellular state in which very few cells or, indeed, none at all may exist (Altschuler & Wu, 2010). Advances in areas such as cell isolation using microfluidics or microwell devices, preparation of NGS libraries from ultra-low quantities of nucleic acid and innovative labelling strategies for proteomic mass spectrometry have enabled the characterisation of DNA (Navin et al., 2011), RNA (Tang et al., 2009) and proteins (Budnik, Levy, Harmange, & Slavov, 2018) from individual cells. In particular, transcriptomic analysis of single cells has matured rapidly and the technique is now cost effective, highly accurate and capable of determining the distribution of expression levels in tens of thousands of single cells simultaneously (Fan, Fu, & Fodor, 2015; Klein et al., 2015). These innovations in analytical capability have led to significant advances in many areas of cell biology and have resulted in the identification of rare cell types (Montoro et al., 2018), the discovery of gene expression changes between different cell types (Tirosh et al., 2016) and the improvement of our understanding of fundamental biology processes such as cellular differentiation (Trapnell et al., 2014).

Overcoming the inherent heterogeneity of biological systems to produce consistent and safe medicines is a critical challenge faced by the biopharmaceutical industry. The development of a commercial therapeutic protein manufacturing process begins with the generation of a CHO cell line that is not only highly productive but also exhibits stable performance over extended culture. To limit heterogeneity, cells transfected with the product transgene undergo one or more rounds of single cell cloning with the best clones brought forward in the process. It is accepted, however, that variability in genotype and phenotype cannot be completely eliminated by cloning and that the population inevitably diverges from the parental clone during subsequent growth and cell culture (Frye et al., 2016). The emergence of heterogeneity coupled with the inherent genomic plasticity of CHO cells can lead to unpredictable gradual or sudden reductions in productivity. A diverse array of mechanisms including copy number loss, transcriptional silencing, chromosomal rearrangements and cell stress have so far been shown to cause production instability (Dahodwala & Lee, 2019). CHO cell line instability has the potential to increase time to market and affect regulatory approval of recombinant therapeutic proteins, it is therefore essential that we continue to refine our understanding of this complex phenotype.

In this manuscript, we report application of single cell RNA sequencing (scRNA-seq) for the characterisation of a recombinant CHO cell line. We utilised the technology to acquire > 3,800 gene expression profiles from a clonally-derived cell line that underwent production instability. Through this analysis we observed intra-cell line heterogeneity as well as significantly lower heavy chain transcript levels across the population. Using trajectory analysis, we were able to reconstitute the evolution of the cell line, demonstrate that the emergence of transcriptome heterogeneity is associated with a reduction in the levels of both mAb transcripts and identify CHO cell gene changes associated with production instability.

## Results

### Production instability in the CHO DP-12N1 cell line

The CHO DP-12 cell line [ATCC clone #1934] is an anti-IL8 IgG1 mAb expressing, clonally-derived cell line generated following random transfection of the transgene plasmid (Figure 1a). During the cell line development process the parental clone was selected for expansion from a panel of 20 clones based on the comparatively high titre of 250 mg/ml observed (“United States Patent: 6025158,” n.d.). After an initial period of effective production in our laboratory, anti-IL8 mAb levels diminished dramatically and could no longer be detected consistently at either 72 hrs (Day 3) or 240 hrs (Day 10) using western blotting (Figure S1) or SEC-MS. Further analysis did however confirm the presence of light chain in both the lysate (Figure 1b, Figure S2) and supernatant (Figure 1c, Figure S3) at both time points. In contrast, the heavy chain was only detected in the supernatant at day 10 of culture (Figure 1c, Figure S3). We also performed western blotting for the light chain in the cell lysate and supernatant under non-reducing conditions and detected a potential light chain dimer at both time points (Figure 1d, Figure S4). To confidently identify the LC dimer, we utilised size exclusion chromatography followed by mass spectrometry to analyse the supernatant at day 10. Deconvolution of the resulting spectrum enabled the detection of a single main peak at 47,903.09 Da (Figure 1e) corresponding to the light chain dimer with a single disulphide bond connecting the two monomers (Δ from the theoretical average mass was 0.6 ppm). No species equivalent to the heavy chain or intact monoclonal antibody were identified and the minor peaks present in the spectrum were unable to be characterised due to the complex nature of the sample. For clarity, we refer to our non-producing variant of the cell line as CHO DP-12N1 in this manuscript.

**Figure 1:**
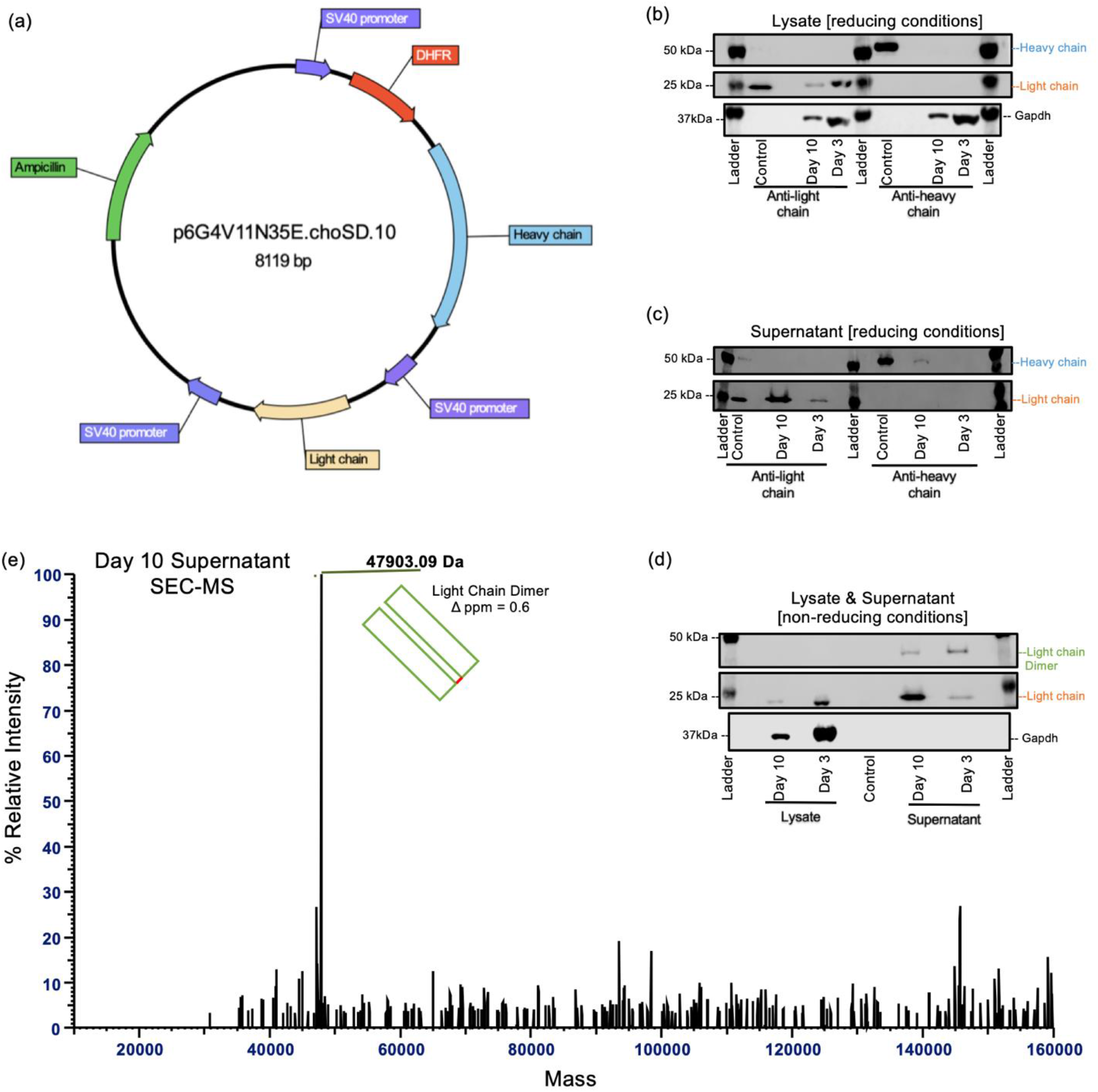
Western blot and mass spectrometry reveals the presence of light chain dimers in the CHO DP-12N1 cell culture media. **(a)** The expression construct for the anti-IL8 mAb was designed to express the heavy and light chains of the antibody under the control of two separate SV40 promoters. Upon confirming that complete anti-IL8 monoclonal antibody could not be detected by either western blot or by mass spectrometry, we analysed the heavy and light chain of the mAb in the cell lysate and the supernatant at Day 3 and Day 10 of cell culture. Under reducing conditions, **(b)** the light chain was detected at both day 3 and day 10 in the lysate, however the heavy chain could not be detected. The heavy chain was detected **(c)** only in the supernatant harvested after 10 days of culture (Day 10). When non-reducing conditions **(d)** were used for western blotting a light chain dimer was present in the supernatant harvested after 3 (Day 3) and 10 days (Day 10) of culture. (0.125 µg per lane of a recombinant human IgG1 kappa (Biorad, HCA192) was used as a control antibody). Mass spectrometry analysis following SEC separation of the supernatant at Day 10 was used to **(e)** confirm the presence of light chain dimer containing one inter molecular disulphide bond.

### Single cell RNA-seq analysis of the CHO DP-12N1 cell line

In this study, we sought to utilise scRNA-seq to examine the CHO DP-12N1 cell line and assess utility of the approach to enhance the study of CHO cell biology through understanding transcriptional heterogeneity and its impact on biopharmaceutical manufacturing. CHO DP-12N1 cells were seeded at 2 × 10^5^ cells/ml in four replicate shake flasks containing chemically defined media (Figure 2a). The cultures reached an average cell density of 2.46 × 10^6^ cells/ml after 72 hrs (> 98% viability for all replicates), at which point samples were acquired for transcriptomic analysis. To compare transcriptomic profiles between aggregated scRNA-seq data and traditional bulk total RNA-seq, a parallel analysis using both approaches was carried out for the four shake flask cultures (Figure 2a).

**Figure 2:**
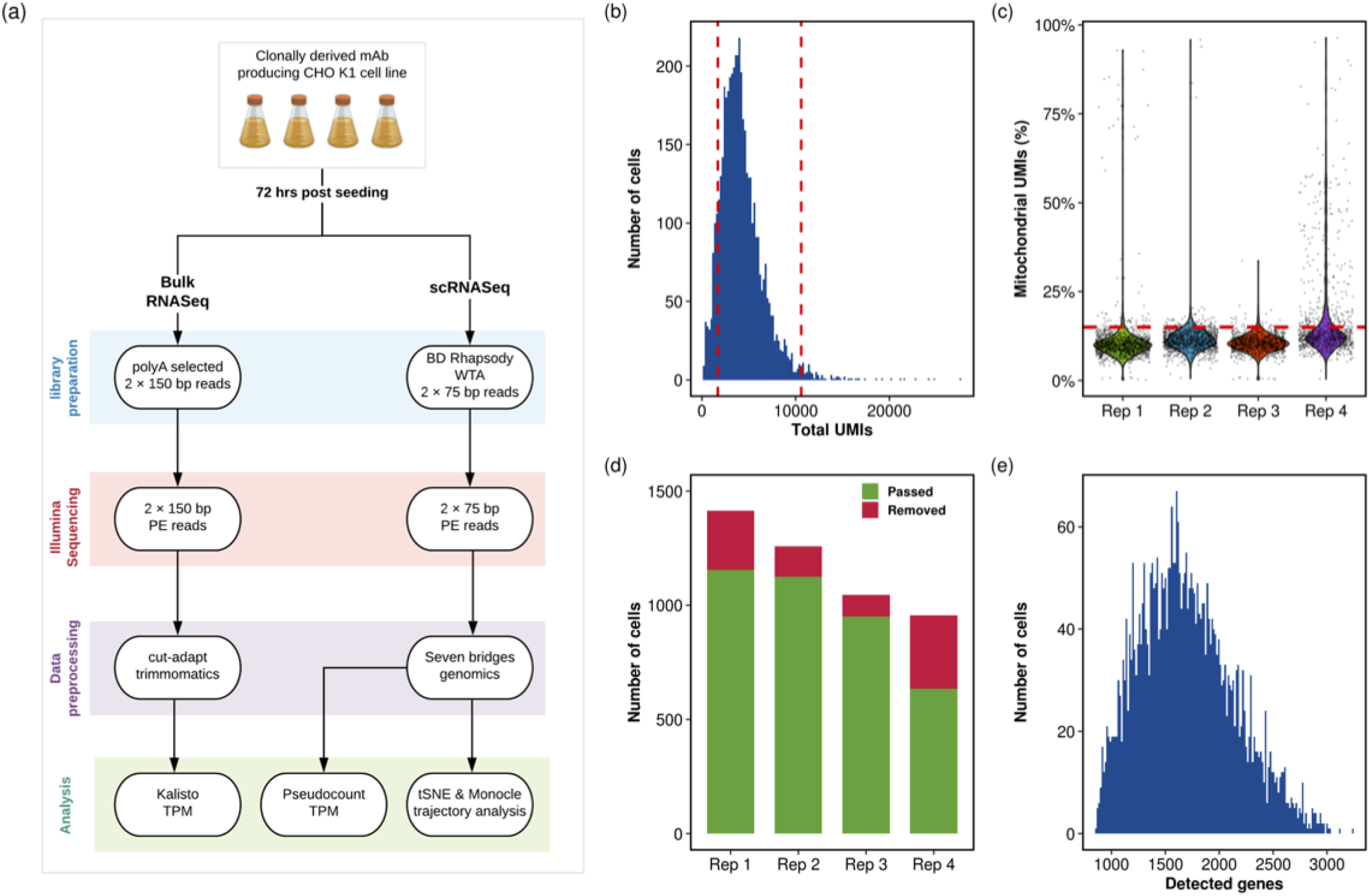
Single cell RNA sequencing using the BD Rhapsody platform enables the capture of thousands of CHO cell gene expression profiles. **(a)** Four biological replicates of the CHO DP-12N1 cell line were cultured for 72 hrs before parallel single cell and traditional bulk RNA-seq analysis were carried out. For the scRNA-seq data, barcodes that were likely from dead or non-viable cells, cells with damaged membranes or more than one cell in a well were eliminated if **(b)** the total UMI count was outside 2.5 × the mean UMI count of the population or **(c)** contained ≥ 15% of UMIs originating from the mitochondrial genes. Following barcode filtering a total of **(d)** 3,866 cells were retained. (**e)** Genes with no detectable expression in more than 100 cells and those genes that were not detected with > 4 UMIs in at least one cell were removed, leaving 2,583 genes for downstream analysis.

For scRNA-seq, ∼20,000 cells from each replicate culture were loaded onto four BD Rhapsody cartridges with ∼200,000 microwells. Beads with oligonucleotide barcodes were added to the cartridge in excess so that virtually all microwells were populated with a bead. Following cell lysis within the microwells, beads and hybridised poly-adenylated RNA molecules corresponding to ∼1,600 captured cells per replicate were transferred into a single tube for reverse transcription. Upon cDNA synthesis, each cDNA molecule was tagged on the 5′ end (the 3′ end of a mRNA transcript) with a molecular index and cell label indicating its cell of origin. The beads were then subject to second strand cDNA synthesis, adaptor ligation, and universal amplification with 22 cycles of PCR. Random priming of the whole transcriptome amplification products to enrich the 3’ end of transcripts was used to prepare sequencing libraries.

Following Illumina sequencing, an average of 86 million 75 bp paired-end reads were acquired for each sample (Table S1). The BD Rhapsody analysis pipeline was used to process sequencing data, conduct sequencing data quality control, and demultiplex cellular barcodes (Figure 2a). Following the completion of this initial pre-processing stage, ∼9% of the sequenced RNA-seq reads were removed from further analysis due to insufficient read length (R1 < 60 or R2 < 42), low base quality (R1 or R2 mean base quality Q < 20) or due to high single nucleotide frequency (R1 ≥ 0.55 or R2 ≥ 0.80), leaving an average of ∼78 million valid reads per replicate (Table S1). 83% of the R1 reads that passed quality control were successfully assigned to cell barcodes following demultiplexing. Mapping of the corresponding R2 reads to the reference genome resulted in a unique alignment rate of ∼76% with ∼43% of these reads mapping against the Ensembl annotated protein coding transcriptome. Upon collapsing to UMIs and application of the RSEC algorithm, between 955 and 1415 unique cell barcodes were identified for the 4 replicate samples and a total of 31 million mRNA molecules detected. The mean numbers of reads and mRNA molecules detected per cell in this experiment were ∼20,320 and 4,276 respectively with an average of ∼1,614 genes detected in each cell (Table S1).

A “pseudo-bulk” gene expression profile (Ding et al., 2020) was generated to enable the comparison of replicate scRNA-seq samples. To construct a pseudobulk expression profile, the UMI counts were summed for each cell barcode captured on the four BD Rhapsody cartridges. The total UMIs from each cartridge were divided by 10^6^ to yield transcripts per million (TPM) expression values for all protein coding genes annotated in the reference genome. The Pearson’s correlation coefficient (PCC) of the log(TPM +1) was calculated between each pair of pseudo-bulk samples with all scRNA-seq replicates found to be highly similar (R^2^ > 0.98) (Figure S5). The matched bulk RNA-seq data was pre-processed and quantified with Kallisto using the same reference genome as the scRNA-seq data to generate gene-level TPM expression levels. The replicate bulk samples were found to be highly similar with each other (Figure S6) and the calculation of the PCC between each scRNA-seq pseudobulk sample and the bulk RNA-seq data from the same sample yielded an R^2^ > 0.8 for all replicates (Figure S7).

For the next stage of our analysis the four raw scRNA-seq cell count tables produced by the BD Rhapsody bioinformatics pipeline were merged to produce a single UMI count matrix comprised of 4,673 cell barcodes and 20,594 genes. To ensure only high-quality gene expression profiles were retained for further analysis, the UMI count matrix was first filtered to remove data that might have originated from non-viable cells, cells with damaged membranes or multiplets (the capture of two or more cells in a single well). To this end, the total number of UMIs captured for each cell barcode was compared to the average number of UMIs detected for the population. 522 cell barcodes with a UMI count that was beyond 2.5 times the mean UMI count (Figure 2b) were removed. A further 285 cell barcodes with > 15% of detected UMIs originating from the mitochondrial genome (Figure 2c) were eliminated. In addition to removing poor quality cell barcodes, we also filtered genes from the UMI count matrix. Only those genes that were detected in more than 100 cells and had a maximum expression > 4 UMIs in at least one cell were retained. Finally, the 13 protein coding genes encoded in the mitochondrial genome were excluded from downstream analyses. The final gene expression matrix was comprised of 3,866 high quality cells with UMI counts for 2,583 genes (Figure 2d & Figure 2e).

### Identification of intra-population differences in transcription in the CHO DP-12N1 cell line

To conduct an exploratory analysis of CHO DP-12N1 scRNA-seq data we utilised the Monocle v2 R package (Qiu et al., 2017; Trapnell et al., 2014) to perform an unsupervised dimensionality reduction and visualisation of the 3,866 cells using t-distributed stochastic neighbour embedding (t-SNE) (Maaten & Hinton, 2008). For the first stage of this analysis, 2,545 variable genes were selected from the UMI count matrix (Figure 3a). The dimensionality of the single cell expression data was further reduced using principal components analysis (PCA) and following determination of the proportion of variance captured by each component, the first 12 PCs were selected for t-SNE (Figure 3b).

**Figure 3:**
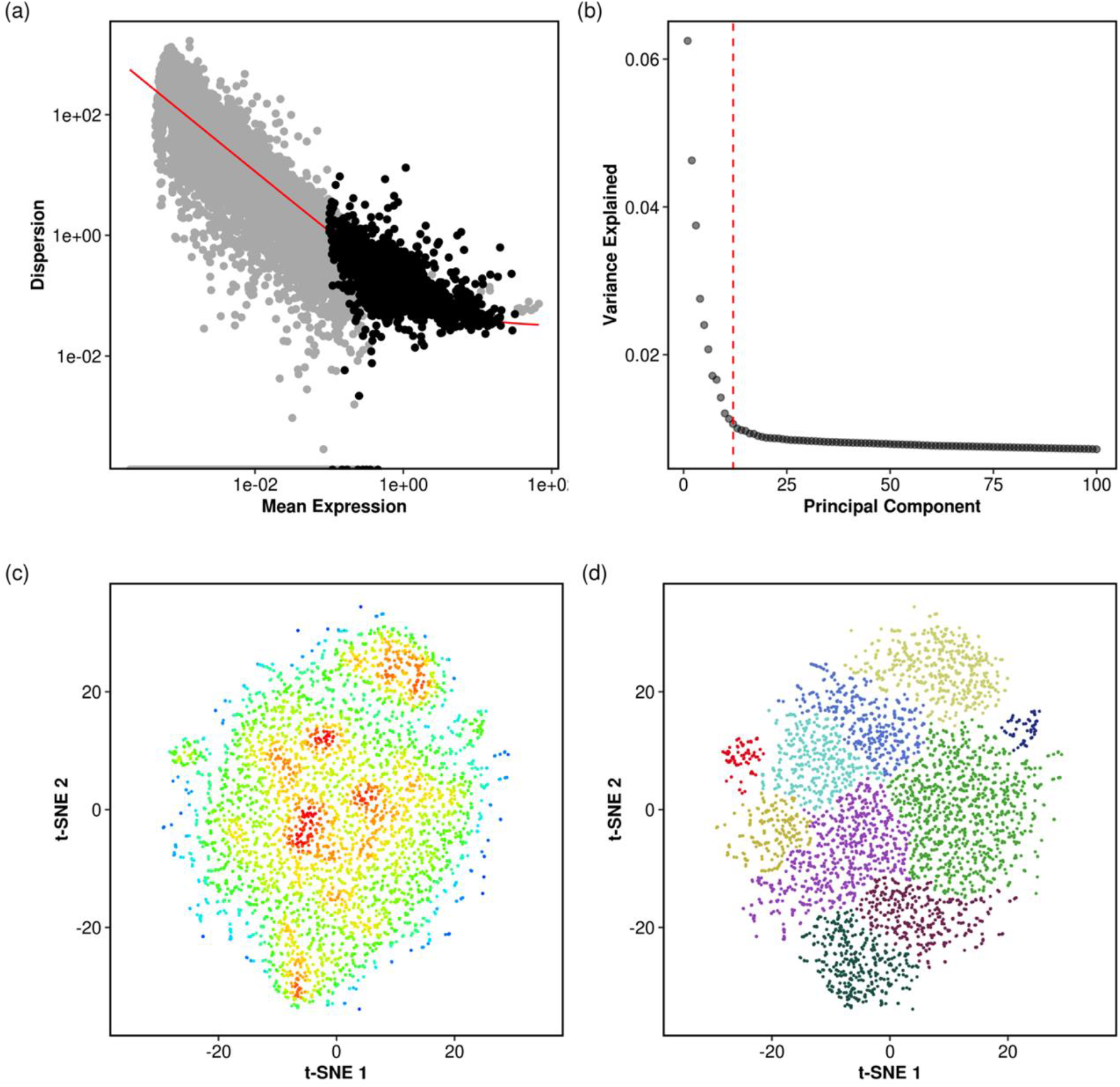
Single cell transcriptomics reveals intra-population heterogeneity of the CHO DP-12N1 cell line. t-SNE was utilised to obtain a global visualisation of the scRNA-seq data. An initial dimension reduction step was conducted using principal components analysis of **(a)** 2,545 of the most variable genes. The first **(b)** 12 principal components were utilised for t-SNE. Plotting (**c)** the first two t-SNE components and determining the local density of cells in the two-dimensional space revealed the presence of regions of higher density. **(d)** Utilisation of the Monocle densityPeak clustering algorithm identified 10 clusters from the t-SNE analysis.

To account for potential heterogeneity from technical and biological confounding factors, we corrected the UMI count data prior to t-SNE, ensuring that covariation between multiple factors was removed simultaneously. For technical covariates, possible batch effects arising from use of different cell isolation cartridges as well as differences in the total number of UMIs captured and genes detected per cell were regressed out. The most common biological covariate, cell cycle, was also eliminated using the Cell Cycle Scoring method in the Seurat package (Butler, Hoffman, Smibert, Papalexi, & Satija, 2018). To utilise this method, the predefined genesets for the S and G2/M phases of cell cycle provided by Seurat were mapped to Chinese hamster genes. The expression levels of these genes were used to assign a score to each cell indicating the likelihood of a given cell being in either S or G2/M phase. For the 3,866 cells, 1,484 and 1,579 were classified as being in S or G2/M respectively while the remaining 803 cells were designated as G1. Using the assigned scores, cell cycle variation was regressed out prior to t-SNE dimensionality reduction. Successful correction of the data was confirmed following visualisation of a uniform distribution of cells from each cell cycle phase across the t-SNE space (Figure S8).

No dramatically distinct clusters of cells were observed from the two-dimensional t-SNE plot (Figure 3c). Visualisation of the density of points in t-SNE did however, reveal several sub-clusters of cells confirming the presence of transcriptional heterogeneity in the CHO DP-12N1 cell line. The Monocle densityPeak algorithm (rho and delta parameters set at 2 and 4 respectively (Figure S9)) was used to identify clusters from the two-dimensional t-SNE representation of the 3,866 gene expression profiles, resulting in the identification of 10 clusters ranging from 57 to 932 cells (Figure 3d, Figure S10).

### Transcriptional heterogeneity in the CHO DP-12N1 cell line is associated with differences in anti-IL8 mAb gene expression in individual cells

For the next stage of our analysis, we investigated the possibility that variability in the expression of the genes encoding the recombinant protein could be a factor in the transcriptional heterogeneity observed within the CHO DP-12N1 cell line. The anti-IL8 mAb is produced via a dicistronic vector expressing the heavy and light chains of the mAb with dihydrofolate reductase (*DHFR*) as a selection marker (“United States Patent: 6025158,” n.d.). The expression of *DHFR* and the heavy chain are controlled by a SV40 promoter while the light chain is controlled via a second, distinct, SV40 promoter (Figure 1a). To enable quantitation of the heavy and light chain genes of the anti-IL8 mAb the transgene sequence was included in the reference genome for both the scRNA-seq and bulk RNA-seq data.

From the bulk RNA-seq data we found that both the heavy and light chain were expressed in the top 2% of all genes. The expression of the light chain (median TPM = 7.9) was higher than that of the heavy chain (median TPM = 6.4) (Figure S11). Comparison of the relative expression of the heavy and light chains across the 3,866 cells analysed by scRNA-seq, similar to the bulk RNA-seq, showed that light chain gene expression was significantly higher across the population (Figure 4a). We also assessed the relative expression of both genes encoding the mAb within the clusters of cells identified from the t-SNE analysis and observed significant differences in transcription of the heavy (Figure 4c) and light chain (Figure 4d) amongst the groups. To visualise the distribution of mAb gene expression across the population, we superimposed the relative expression of the heavy (Figure 4e) and light chain genes (Figure 4f) onto the tSNE plot. The light chain was expressed not only at a higher level but also more consistently across the population in comparison to the heavy chain. We found that ∼14% of the cells captured (n = 566) had no detectable heavy chain expression, while in contrast only 5% of cells (n=208) had no detectable light chain expression.

**Figure 4:**
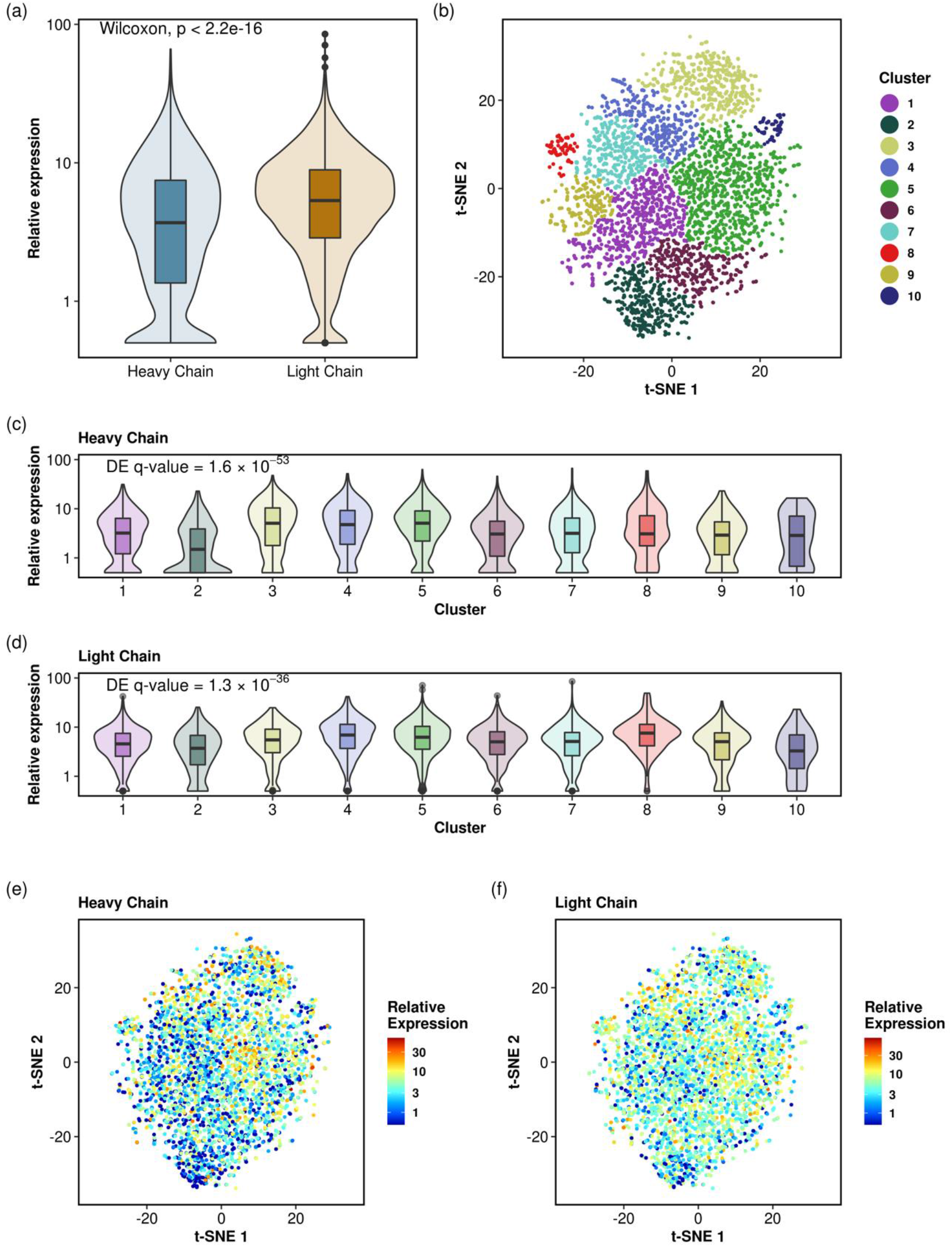
Expression of anti-IL8 mAb heavy and light chain mRNA is a factor in transcriptional heterogeneity of the CHO DP-12N1 cell line. **(a)** The proportion of light chain mRNA expressed is significantly higher than that of the heavy chain across the population. Significant differences for **(b)** the intra cell line clusters for the **(c)** heavy and **(d)** light chain. The q-value was calculated using the Monocle *differentialGeneTest* function while correcting for biological and technical covariates. These difference in expression across the population are reflected by the distribution of cells in the t-SNE space plot where the **(e)** heavy chain is observed to be expressed at a lower level toward the bottom half of the plot while **(f)** the light chain is expressed more uniformly across the cell population.

### Pseudo-temporal ordering of the CHO DP-12N1 scRNA-seq data resolves a single cell trajectory correlated with a decrease in anti-IL8 mAb gene expression

To understand the emergence of heterogeneity and evolution of the cell line we applied a technique known as pseudotemporal ordering or trajectory analysis (Trapnell et al., 2014). Briefly, trajectory analysis is an unsupervised approach that enables the identification of routes (a trajectory) through cellular space that minimise the distance between similar cells. Each cell can be ordered along the trajectory and the distance from the beginning or “root” of the trajectory determined with a measure termed “pseudotime”. To conduct this analysis, 1,322 genes that were differentially expressed between the 10 clusters identified from the t-SNE analysis were used as input to Monocle DDRTree algorithm (Qiu et al., 2017; Trapnell et al., 2014) and the data were again corrected for potential confounding technical and biological factors. Trajectory analysis identified 9 distinct groups or “states” for the CHO DP-12N1 cell line (Figure 5a) ranging from 50 to 1007 cells (Figure 5b).

**Figure 5:**
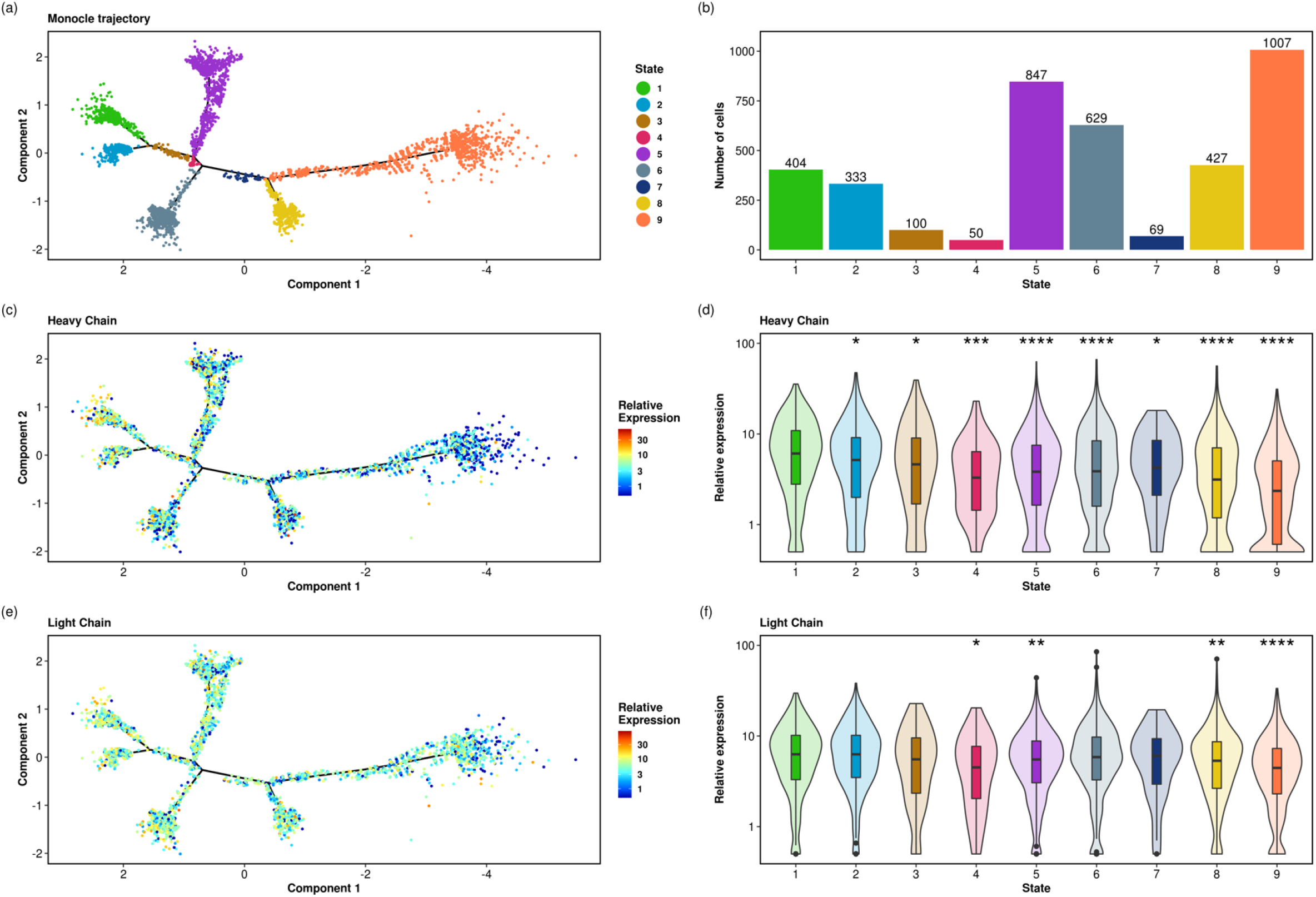
The CHO DP-12N1 trajectory captures the decrease in anti-IL8 mAb gene expression. **(a)** To construct a trajectory using the Monocle DDRTree algorithm, 1,322 genes differentially expressed between the t-SNE clusters were identified before the DDRTree algorithm was utilised to produce the single cell trajectory. **(b)** The algorithm identified 9 distinct cell states ranging from 1007 to 50 cells. To determine if there was a difference in transcription of the heavy and light chains across the trajectory and to identify the root state, we determined the relative expression in each of the nine cell states identified by the monocle DDRTree algorithm. **(d)** The cells in cell state 1 had significantly higher transcription of the heavy chain when compared to all other states, while **(f)** the light chain remained consistent for longer across the trajectory. The pattern of the **(c)** heavy and **(e)** light chain is evident when the expression values are overlaid on the trajectory plot. Note: the x-axis of **(a), (c)** & **(e)** is reversed for clarity (* p < 0.05; ** p < 0.01; *** p < 0.001).

To identify the root state of the trajectory we assumed that high transcription of both mRNAs was essential to the anti-IL8 titre of the original clone and that the cell state identified in this study with the highest heavy and light chain gene expression would most closely resemble the original clone. To assess the expression levels of the mAb genes, the 3,866 cells were stratified based on cell state. For the heavy chain, cell state 1 was found to have a significantly higher median relative expression in comparison to other cell states with cell state 9 having the lowest median relative expression (Figure 5c). The highest light chain expression was observed for cell state 1 although the difference between cell states 2, 3 6, and 7 was not statistically significant (Figure 5e). Cell state 9 again had the lowest relative median expression for the light chain. Upon overlaying the relative expression on the trajectory plot, the decrease in heavy chain expression across the trajectory is evident (Figure 5d). While the light chain expression diminishes toward the end of trajectory (Figure 5e), the decrease is not as pronounced as the heavy chain (Figure 5c). Based on these results, we set the root of the CHO DP-12N1 trajectory at cell state 1 and ordered the pseudotime variable accordingly (Figure 6a).

**Figure 6:**
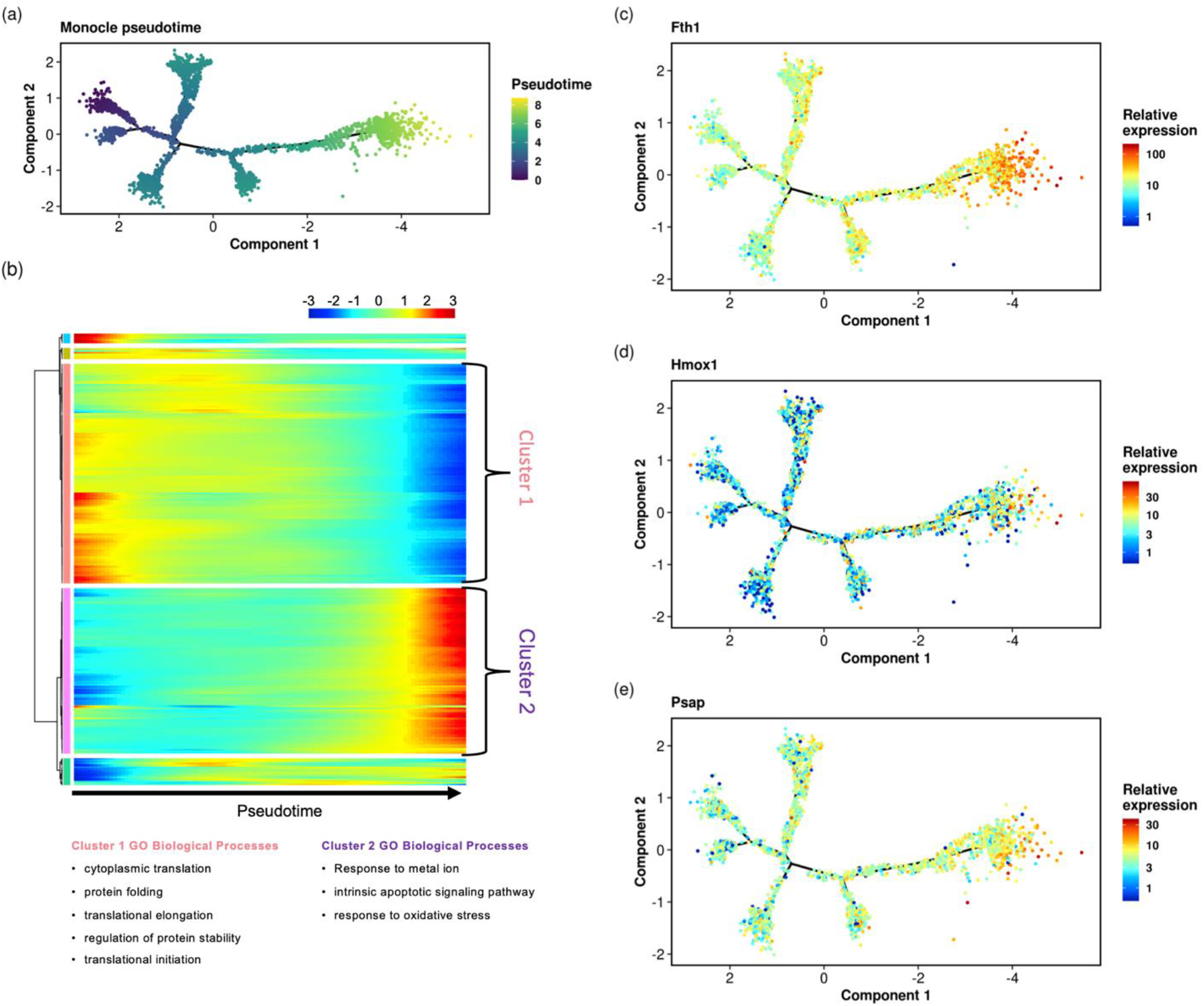
Hierarchical clustering analysis identifies groups of genes that increase or decrease as cells progress and reveals the overrepresentation of cellular processes and alterations in the expression of individual genes. (**a)** following the identification of the cells most similar to the original isolated clone (i.e. those with the highest expression of both heavy and light chain) the pseudotime variable was set to increase as cells diverged, **(b)** cluster analysis partitions genes associated with Pseudotime into 5 groups of genes that increase or decrease in expression. Enrichment analysis against GO of the two largest gene clusters identifies the overrepresentation of biological processes including protein synthesis, apoptosis and the response to oxidative stress. Individual genes related to the identified pathways including **(c)** *Fth1*, **(d)** *Hmox1* and **(e)** *Psap* were found to correlate with progression through the single cell trajectory. Note: the x-axis of **(a), (c), (d)** & **(e)** is reversed for clarity)

### Gene expression changes related to protein synthesis and the response to cell stress are associated with the progression of cells along the CHO DP-12N1 trajectory

In order to understand the transcriptomic changes associated with the progression of cells along the CHO DP-12N1 trajectory, genes that positively or negatively correlated with pseudotime were identified. Potential confounding factors such as cell cycle, batch, total UMI and detected genes per cell were again regressed from the analysis prior to utilising the Monocle differentialGeneTest function, with 880 genes found to be significantly (BH adj. p-value < 0.01) associated with pseudotime (Table S3). The ferritin heavy chain 1 (*Fth1*) gene was found to be the gene that changed most significantly and was dramatically upregulated at the end of the single cell trajectory (Figure 6c). Hierarchical clustering analysis of the differentially expressed genes resulted in the identification of 5 gene clusters (Figure 6b) that followed a similar expression pattern. The two largest clusters were subsequently found to be enriched for multiple biological processes (Table S4). The first cluster, comprised of 446 genes that tended to decrease in expression as cells progressed along the trajectory (and were correlated to a decrease in heavy and light chain expression), was found to be enriched for genes involved in protein synthesis and stability. Biological processes relating to the response to oxidative stress and apoptosis e.g. *Hmox1* (Figure 6d) and *Psap* (Figure 6e) were found to be overrepresented in the second cluster of 336 genes that increased in expression as pseudotime increased.

## Discussion

Although the development of a highly productive CHO cell line begins with a single cell cloning step in order to limit heterogeneity in the cell population, eliminating heterogeneity entirely from a master cell bank is not possible (Frye et al., 2016). Not only are the emergence of mutations and chromosomal rearrangements inherent to the culture of immortalized cells but the exertion of various stresses (e.g. selection and media adaption) as well as epigenetic changes at subsequent stages of cell line and process development have the potential to alter the cell population further. For these reasons, while single cell cloning is recognized as a valuable stage, greater focus is placed on ensuring stability of the manufacturing process and critical quality attributes of the final product. Consequently, long term stability studies are required to ensure a production cell line maintains desirable growth rates and productivity and crucially that critical quality attributes are consistent throughout manufacturing to ensure patient safety (Dahodwala & Lee, 2019). The adoption of new high throughput approaches that enable increased characterisation of cell populations has the potential to provide greater assurance as well as enhance our fundamental understanding of the origins and impact of heterogeneity on the cell line performance.

In this study, we assessed the utility of scRNA-seq to characterise transcriptional heterogeneity in a CHO cell line following the acquisition of > 3,800 gene expression profiles. Biological replicates measured by scRNA-seq were highly similar and indeed aggregated single cell profiles from the same sample, when merged, were comparable to their matched bulk RNA-seq counterpart. Global assessment of the population using t-SNE revealed CHO cell heterogeneity did not result in, as one might expect from a clonally-derived cell line, a dramatic separation of cells as has been reported for complex samples such as tissues. Nevertheless, these data show that transcriptome-wide variation is present in this clonally derived cell line. From the t-SNE and cluster analysis we found that variation in anti-IL8 mAb transgene expression was a factor in the distribution of cells in the 2-D space, reflecting the burden exerted on cells through the production of a recombinant protein and the interplay between transcriptome heterogeneity and transgene expression.

The CHO DP-12 cell line is prone to production instability (Beckmann et al., 2012) and over time in our laboratory the production of the anti-IL8 mAb reduced dramatically. The heavy chain transcript of the mAb was expressed at a significantly lower level in comparison to the light chain transcript (correlating with the protein levels detected at Day 3 and Day 10). We also confirmed the presence of a light chain dimer in the supernatant of the CHO DP-12N1 cell line. Approximately 14% of the cells analysed in this study had no detectable heavy chain transcripts, consistent with the excess of light chain proteins observed. Our findings are in agreement with an independent study of production instability in the CHO DP-12 cell line over long term culture which also reported the loss of the heavy chain expression (Beckmann et al., 2012) and a light chain only subpopulation of cells.

Using scRNA-seq data and trajectory analysis we were also able to trace the evolution of the CHO DP-12N1 cell line over its lifetime in culture to gain an understanding of CHO DP-12N1 transcriptome heterogeneity and the cellular processes associated with production instability. For this analysis we considered the emergence of heterogeneity in a clonally-derived cell line as a gradual, continuous process spanning a range of cell states that began with cells that were most similar to the original isolated clone and continued to the most diverged. Trajectory analysis enabled the 3,866 cells to be separated into 9 distinct cellular states and ordered along a path spanning transcriptomic space. We chose not to focus on the branching points in this study, as a complete history of the cell bank was unavailable. For future studies, cells could be analysed at different timepoints throughout the cell line development process and the differential expression between any branch points could be determined (i.e. after selection or expansion). It is important to note that, as with the t-SNE analysis, we resolved the trajectory in a purely data driven, unsupervised manner. We did, however, utilise basic knowledge of the CHO DP-12N1 cell line by identifying the cell state with the highest expression of both heavy and light chain and we assumed that these cells were most similar to the isolated clone from which the cell line was expanded to determine the start of the trajectory. Overlaying the relative expression of the heavy and light chain for each cell along the trajectory confirmed that divergence from the beginning of trajectory was correlated with decrease in expression of both genes. The heavy chain expression was significantly reduced at an earlier point in the trajectory compared to the light chain which could also contribute to production instability and/or the formation of light chain dimers.

To examine the underlying biological mechanisms associated with the emergence of heterogeneity and production instability in the CHO DP-12N1, we identified 2 distinct groups of CHO cell genes that tend to increase or decrease as cells progressed along the trajectory. We found that the group of genes that decreased in expression were involved in protein synthesis, protein folding and stability. As might be expected as mAb heavy and light chain transcript levels decrease, there is a lower demand on the protein synthesis machinery to produce the recombinant protein, which could explain the observed reduction in genes involved in protein production and assembly. As cells progressed along the trajectory, genes involved in apoptotic signalling pathways and in response to oxidative stress and metal ions were found to increase in expression. Amongst the upregulated genes involved in oxidative stress response were *Hmox1, Fth1* and *Psap*. Psap has been shown to protect neural cells against oxidative stress (Ochiai et al., 2008). The upregulated *Hmox1* gene encodes heme oxygenase 1, an enzyme that is predominantly controlled at the transcriptional level and is upregulated in response to oxidative stress (Gozzelino, Jeney, & Soares, 2010). Hmox1 converts heme to CO, Fe^+2^ and the antioxidant biliverdin (Kerins & Ooi, 2018), which is further metabolised to bilirubin. FTH1 encodes the heavy subunit of ferritin, which exerts its antioxidant role by storing and sequestering iron, produced from the reaction catalysed by Hmox1 (Fraser, Midwinter, Berger, & Stocker, 2011). The increase in the *Fth1, Hmox1* and *Psap* genes could be a cytoprotective mechanism activated in response to reactive oxygen species associated with ER stress (Cao & Kaufman, 2014) induced by differences in the expression of the heavy and light chains of the mAb.

This study demonstrates that scRNA-seq permits the study of CHO cell biology at unprecedented resolution. It is important to note that utilising the method presents a number of challenges when compared to traditional bulk RNA-seq. The low starting amount of mRNA per cell increases technical noise and produces a larger proportion of zero gene expression values than bulk RNA-seq (Hicks, Townes, Teng, & Irizarry, 2018). While a component of these zero values are biological (a particular gene is not expressed in a given cell), only a fraction of each cell’s total transcripts will be present in the sequencing library due to the low capture efficiency of scRNA-seq methods (Liu & Trapnell, 2016). This limitation impacts the computational analysis and interpretation of the resulting data and can prohibit reliable quantitation of very low abundance transcripts. There are however experimental methods available that enable targeted amplification of a panel of specific genes enabling detection of low abundance genes and decrease the required sequencing depth in comparison to WTA scRNA-seq protocols (Mair et al., 2020). The development of targeted gene panels for studying biological processes (e.g glycosylation) and/or regulatory genes essential for the efficient manufacture of recombinant proteins could be valuable in the future for scRNA-seq analyses in CHO cells.

In summary, scRNA-seq is a powerful tool for understanding how heterogeneity in recombinant therapeutic protein producing CHO cell lines impacts process performance. The application of single cell transcriptomics has enabled the study of transgene and host cell gene expression patterns at unprecedented resolution. We were able to trace the evolution of a cell line, capturing the divergence of the population over its life time in culture and the associated change in transgene expression. We anticipate that further application of scRNA-seq has the potential to dramatically improve our knowledge of CHO cell biology and enable more precise genetic engineering targets to increase the efficiency of the therapeutic protein manufacture.

## Methods

### Cell Culture

The CHO cell line utilised for this study was originally acquired from the ATCC CHO DP-12 cells [clone#1934 aIL8.92 NB 28605/14] (ATCC CRL-12445™). In this paper, we refer to the CHO DP-12N1 to differentiate the non-producing cell line in our laboratory from the reference cell line available through ATCC. CHO DP-12N1 cells were cultivated in BalanCD CHO Growth A medium (Irvine Scientific, 91128-1L) supplemented with 6.7 mM L-glutamine in a total volume of 30 mL in 250 mL flasks (PC Erlenmeyer Flasks with vented caps, Fisher 100022611). Methotrexate (300 nM; Sigma-Aldrich M9929) was added to the culture medium every third passage to maintain stability of anti-IL8 gene expression. Incubator conditions were set to 37°C, 5% CO_2_ and a relative humidity of 70% with a shaker revolution of 170 rpm (Troemner Talboys Advanced Dura Shaker, NC1400130, exterior microprocessor controller). Cells were passaged every three to four days with density not exceeding 8.5 × 10^6^ cell/ml. Manual Trypan Blue exclusion (0.4%) was used for cell counting and viability determination on an inverted bright field microscope.

For bulk and single cell RNA-seq four biological replicates (n=4) were seeded at 2 × 10^5^cell/ml and cultured for 72 hours. Cells were pelleted by spinning at 1000 × g for five minutes. Supernatant was removed and the cells were resuspended in freezing medium (Embryomax Cell Culture Freezing Media with DMSO (EMD Millipore, S-002-D) at a final concentration of 1 × 10^7^cells/ml/cryovial. Cells were stored in Mr. Frosty (Nalgene®, C1562) at -80°C overnight and then transferred to liquid nitrogen (−130°C).

For western blotting and mass spectrometry CHO DP-12N1 cells were cultured in Mini Bioreactor Tubes (Corning™ 431720) in 5 ml working cultures at the seeding density and cell culture conditions described above. The supernatant and cells were harvested after 3 or 10 days of culture.

### Western blotting

Culture medium was separated from the cells following a 5-minute centrifugation at 1,000 g. Cells were lysed in RIPA buffer (Thermo Scientific™ cat.no. 89900) supplemented with 1% Halt protease inhibitor cocktail (Thermo Scientific™ cat.no. 87786). SDS-PAGE was performed using 4-12% Bolt™ Bis-Tris gels (Life Technologies, NW04127BOX). Separated proteins were blotted onto nitrocellulose membranes (pore size: 0.45 μm) by the Invitrogen Power Blotter–Semi-dry Transfer System (Thermo Scientific™) following a 12 minutes transfer. Membranes were blocked at room temperature for 1 hour with the Odyssey® Blocking Buffer (Licor, cat.no. 927-50000) before incubation with primary antibodies overnight at 4°C. The heavy chain was detected by the anti-human IgG Fcγ fragment specific primary antibody (1:1,000; Jackson, cat.no.109-005-008), the light chain by the anti-human kappa light chain antibody (1:2,500; Biorad, cat.no. STAR 127) and GAPDH by the anti-GAPDH monoclonal antibody (1:10,000; Proteintech, cat.no. 60004-1-Ig). Anti-goat IR-Dye (Licor, cat.no. 926-68074) and anti-mouse (Licor, cat.no. 926-32210) secondary antibodies were used. All primary and secondary antibodies used were diluted in 5% non-fat milk in PBS. The Precision Plus Protein™ Dual Color Standards (Biorad, cat.no.1610374) was used for molecular weight estimation. The recombinant Human IgG1 Kappa (Biorad, cat.no. HCA192) was used as a positive control. For samples resolved under reducing conditions DTT was added to a final concentration of 50mM.

### Mass spectrometry

For size exclusion chromatography-mass spectrometry (SEC-MS) supernatant collected at 240 hrs was filtered with 0.45 µm and 0.2 µm low-binding protein Durapore^®^ filters (Merck Millipore). 25 µL of clarified supernatant was directly injected on a Vanquish™ uHPLC (Thermo Sceintific) equipped with a MAbPac™ SEC-1 4.0 × 300 mm column (Thermo Scientific) hyphenated to an Exploris 240 Orbitrap mass spectrometer (Thermo Scientific). SEC separation was carried using a 0.300 mL/min flow rate of 50mM ammonium acetate (Merck Sigma) for 20 minutes in isocratic conditions. Column oven was maintained at 30°C. MS acquisition was performed through Chromeleon CDS 7.2.10 software and MS settings were as follows. *Source settings*: Spray voltage in positive mode at 3.8 kV, sheath gas 25 arbitrary units (au), auxiliary gas 10 au, ion transfer tube 300°C, vaporizer temperature 225°C. *Scan properties*: scan range 2,500-8,000 m/z, RF lens 60%, acquisition gain control 300%, max injection time 200 ms, 10 microscans, in-source fragmentation on use 80V, Orbitrap resolution 30,000 (at m/z 200). Application mode was intact protein, using high pressure mode for HCD cell. Raw data were analysed using BioPharma Finder™ v. 3.2 software (Thermo Scientific) using the ReSpect™ algorithm with the “average over selected time” range function with the following settings: mass range 2,500-8,000 m/z, output mass range 10,000 – 160,000 Da, charge states deconvolution tolerance 20 ppm, charge states between 10 and 50, minimum adjacent charge states 3.

### Bulk RNA-seq library preparation

Cells frozen in freezing medium (Embryomax Cell Culture Freezing Media with DMSO (EMD Millipore, S-002-D)) and stored in liquid nitrogen for 5 days were thawed and pelleted by centrifugation and the supernatant was discarded. The cell pellet was resuspended in 1 ml Trizol (Invitrogen, cat. no 15596026) and stored at -80°C until used. Total RNA was isolated from cells using Trizol (Invitrogen, cat. no 15596026) following manufacturer instructions. For each sample, 2.5 μg of total RNA were poly(A) enriched with the Seq-Star™ poly(A) mRNA Isolation Kit (cat. no AS-MB-006) and RNA libraries were prepared using the KAPA Stranded RNA-seq library preparation kit (Kapa Biosystems, cat. no KK8401) according to manufacturer’s specifications. RNA samples were fragmented for 6 minutes at 85°C prior to library preparation and the final library was amplified with 8 PCR cycles. Libraries were sequenced on an Illumina HiSeq4000 (Illumina, San Diego, CA) configured to yield 40 million 150bp paired-end reads per sample.

### Bulk RNA-seq data analysis

Cutadapt (v.1.18) (Martin, 2011) was used to remove adapters from the raw RNA sequencing reads before quality trimming was performed using Trimmomatic v0.36 (Bolger, Lohse, & Usadel, 2014). STAR v2.7.2d (Dobin et al., 2013) was utilised to align reads to the Ensembl v99 CriGri-PICR (GCA_003668045.1) genome. At present the Cri-Gri PICR genome assembly does not include the mitochondrial genome. A hybrid reference genome was utilised for RNA-seq read mapping using the mtDNA sequence of the CHO-K1 reference genome (GCA_000223135.1). The anti-IL8 mAb expression construct used to generate the CHO DP-12 cell was also added to the reference genome to enable the quantitation of heavy and light chain transcripts. To determine the expression levels of genes the hybrid reference genome was used in conjunction with Kallisto (Bray, Pimentel, Melsted, & Pachter, 2016) to calculate an aggregated transcripts per million (TPM) expression value for each gene.

### Single cell RNA-seq

For single cell gene expression profiling, each cryovial containing a biological replicate was removed from the nitrogen tank, thawed at 37°C, diluted with PBS and centrifuged at 180x g for 5 minutes. The supernatant was removed without disturbing the pellet, gently resuspended and centrifuged at 300x g for 5 minutes. The supernatant was discarded, and the cell pellet was resuspended in 1 mL of cold Rhapsody Sample Buffer and kept on ice. 3.1 μL of 2 mM Calcein AM and 3.1 μL of 0.3 mM Draq7 were added to 620 μL volume of the cell suspension (1:200 dilution). The suspension was gently pipetted up and down to mix well, then incubated in the dark in a heat block at 37°C for 5 minutes. The stained cells were gently mixed by pipette and immediately counted. Ten μL of the cell suspension was pipetted into one chamber of a INCYTO™ disposable hemocytometer, loaded onto the Rhapsody Hemocytometer Adapter and counted in the BD Rhapsody™ Scanner ≤5 minutes after loading. After scanning was complete the results were analyzed using the Rhapsody software to view the total cell concentration and cell viability. These calculations were used to load approximately 20,000 cells into each BD Rhapsody cartridge. After loading, cell count and viability was captured using the Rhapsody imaging system, the cells were lysed, and magnetic beads bound with cellular mRNA were retrieved. A proportion of beads corresponding to 1,600 cells were used to generate libraries according to the “BD Rhapsody™ System mRNA Whole Transcriptome Analysis (WTA) and Sample Tag Library Preparation Protocol. scRNA-seq libraries were sequenced using an Illumina NextSeq 500 (Illumina, San Diego, CA) configured to yield 75 bp paired end reads.

### scRNA-seq data analysis

#### Generation of a UMI count matrix

For pre-processing and demultiplexing of cellular barcodes, the FASTQ files for each replicate were processed by the BD Rhapsody WTA bioinformatics workflow (BD Biosciences) on the Seven Bridges Genomics (SBG) cloud platform using the default parameters. The first phase of the pipeline establishes the quality of each sequenced read pair by assessing read length, mean base quality and single nucleotide frequency (SNF). Read pairs where the first read (R1) was less than 66 nt or the second read of the pair (R2) was less than 64 nt, were eliminated from further analysis. Read pairs were also discarded if the average base quality of either read was < 20 or if SNF was ≥ 0.55 for R1 or ≥ 0.80 for R2. The pre-processed data was then utilised to identify the cell barcode and the unique molecular index (UMI) from the R1 read (the R1 read only contains a sequence identifying the cell). The R1 read contains 3 sequences, each separated by a common sequence, which are combined to produce the final cell barcode and a UMI sequence downstream of the third barcode. Reads were first checked for perfect matches in all 3 barcode sequences and retained, while the remaining reads were subject to a further round of filtering. R1 reads with base substitutions, insertions, deletions and errors were recovered for further processing.

The R2 reads contain sequence information from the mRNA captured. The R2 read was aligned to a STAR index generated from an identical reference genome to that of the bulk RNA-seq (i.e. the Chinese hamster nuclear genome, the CHO-K1 mitochondrial genome and the anti-IL8 plasmid). Upon completion of the STAR mapping phase, UMI sequences were collapsed. To remove the effect of UMI errors on the final molecule count, an adjustment algorithm termed recursive substitution error correction (RSEC) was used -RSEC identifies single base substitution UMI errors and adjusts to the parent UMI sequence. RSEC uses two factors in error correction: 1) similarity in the UMI sequence and 2) raw UMI coverage. Molecules containing UMI sequences that differed by one base were collapsed and the sum of the reads output as “RSEC-adjusted molecules”. RSEC counts were subsequently used for downstream analyses. The final output from the BD Rhapsody WTA bioinformatics workflow was a cell/gene count matrix for each scRNA-seq sample.

#### Filtering the UMI count matrix

The replicate cell/gene matrices outputted from the SBG pipeline were imported into the R v4.0 Statistical Software Environment and merged to form a single matrix for further analysis. Cell barcodes that could have originated from non-viable cells were removed by analysing the number of UMIs detected from each cell before comparison to the average UMIs detected for the population. The proportion of UMIs originating from mtDNA was also determined for each cellular barcode and those with > 15% mitochondrial UMI counts were eliminated from further analysis. Genes that were not detected in any of the cells captured, as well as genes with a UMI count < 4 in any cell were also eliminated.

#### t-SNE, trajectory and differential expression analysis

The reduceDimensonality function in the Monocle 2 R package was utilised to conduct t-distributed stochastic neighbour embedding (t-SNE) as well trajectory analysis using the DDRTree algorithm. The Monocle differentialGeneTest function was used to detect genes that differed between clusters identified following t-SNE as well as those genes associated with cell progression along the trajectory. Those genes with a BH adjusted p-value < 0.01 were considered significant. Potential confounding technical and biological factors were eliminated prior to each analysis.

#### Cell cycle correction

Prior to dimensionality reduction, trajectory analysis and differential gene expression analysis, cell cycle effects were eliminated. For this analysis the cell cycle scoring function in Seurat v3 (Butler et al., 2018) was first used to assign a score indicating the likelihood that each cell was in either the S or G2/M phase. Seurat provides precompiled lists for human and mouse genes known to play a role in distinct phases of the cell cycle. To conduct this procedure, we mapped the murine gene list to the Chinese hamster genome to carry out the classification. The resulting scores for S and G2/M phase were used to regress out the effect of cell cycle in downstream analyses.

#### Enrichment analysis

The overrepresentation of gene ontology (GO) biological processes within gene lists found to have a statistically significant association with pseudotime were identified with the R WebGestaltR package (Liao, Wang, Jaehnig, Shi, & Zhang, 2019). GO biological processes with a Benjamini-Hochberg adjusted p-value of < 0.05 were considered significant.

## Data availability

The scRNA-seq and bulk RNA-seq datasets have been deposited to the Sequence Read Archive (SRA) with accession code PRJNA661407. The raw UMI count matrix for single cell analysis, Kalisto TPM values and code required to reproduce the results presented in this manuscript are available at https://clarke-lab.github.io/CHO_cell_scRNA-seq/

## Author contributions

W.S.D. and C.C. conceived the study and designed experiments; Cell culture, western blots and RNA sequencing were carried by I.T., C.A.S., and N.H; Mass Spectrometry was performed by S.C. and J.B. Data analysis was performed by R.H., and C.C.; C.C., I.T. and R.H. wrote the paper with critical input from N.B. All authors reviewed the paper.

## Acknowledgements

The authors gratefully acknowledge funding from Enterprise Ireland (grant reference: IP/2018/0803/Y), Science Foundation Ireland (grant reference: 15/CDA/3259) and the European Union’s Horizon 2020 framework programme for research and innovation under the Marie Sklodowska-Curie programme (grant agreement No 813453).

## Competing Interests

I.T., S.C., R.H., N.B., J.B., and C.C. declare no competing interests. N.H., C.A.S., and W.S.D. are employees of BD Technologies and Innovation.

## Abbreviations

(CHO): Chinese hamster ovary
(mAb): monoclonal antibody
(NGS): next generation sequencing
(PCC): Pearson’s correlation coefficient
(RNA-seq): RNA sequencing
(scRNA-seq): single cell RNA sequencing
(BH): Benjamini Hochberg
(WTA): whole transcriptome analysis
(UMI): nique molecular index
(MS): Mass spectrometry
(RSEC): recursive substitution error correction
(SEC): Size exclusion chromatography
(SNF): single nucleotide frequency
(t-SNE): t-distributed stochastic neighbour embedding

## Supplementary Tables

**Table S1: Single cell RNA-seq metrics**. The outputs from the Seven Bridges Genomics bioinformatics pipeline are shown including the read pre-processing steps and deconvolution of cell barcodes for each of the four replicates.

**Download:** https://app.box.com/s/y9a3edloqzab81qkfg70uv8ukdaarpqg

**Table S2: Bulk RNA-seq metrics**. The outputs of bulk RNA-seq and Kalisto quantitation for the four bulk RNA-seq replicates matched to the single cell RNA-seq data are shown including the number of reads acquired and those remaining after pre-processing. The number of pseudo-aligned reads and uniquely aligned reads from Kalisto are also shown.

**Download:** https://app.box.com/s/v9qj1eopyl59z27rbhxyfveinusjunah

**Table S3: Genes associated with pseudotime**. Following the identification of the root cell state of the CHO DP-12N1 trajectory, 880 genes were found to be significantly altered over the single cell trajectory. The maximum expression, number of cells expressed as well as the q-value is shown for each gene.

**Download:** https://app.box.com/s/o4rilxlg0u6vn6a7ueb56fzf9lwxenck

**Table S4: Enrichment analysis of pseudotime associated gene clusters**. Following the identification of those genes associated the CHO DP-12N1 trajectory, hierarchical clustering was carried out to identify groups of genes with similar expression patterns. Enrichment analysis was carried out to determine if GO biological processes were overrepresented in the identified clusters. Cluster 1 and 2 were found to be significantly enriched for multiple biological processes. Shown are the GO categories found to be enriched along with the adjusted p-value and the genes leading to the enrichment.

**Download:** https://app.box.com/s/iy426s7280esqa9dyys06jnl04150tdr

## Supplementary Figures

**Figure S1:**
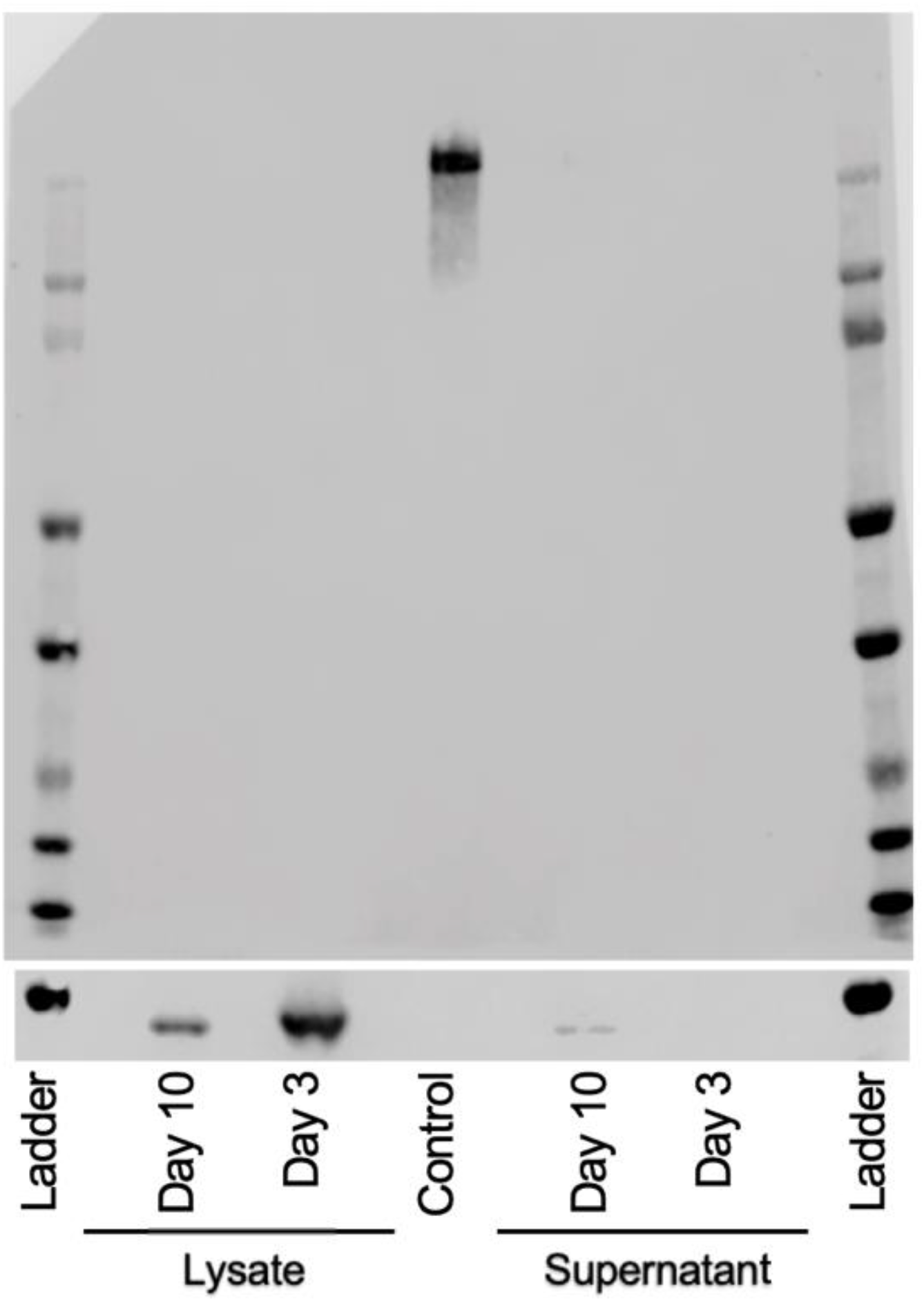
CHO DP-12N1 does not produce intact anti-IL8. Western blot analysis of intracellular and extracellular anti-IL8 antibody under denaturing non-reducing conditions failed to detect the anti-IL8 antibody. Cells and supernatant were harvested after 3 and 10 days of culture and a recombinant human IgG1 kappa antibody (0.125 µg per lane -Biorad, HCA192) was used as a positive control.

**Figure S2:**
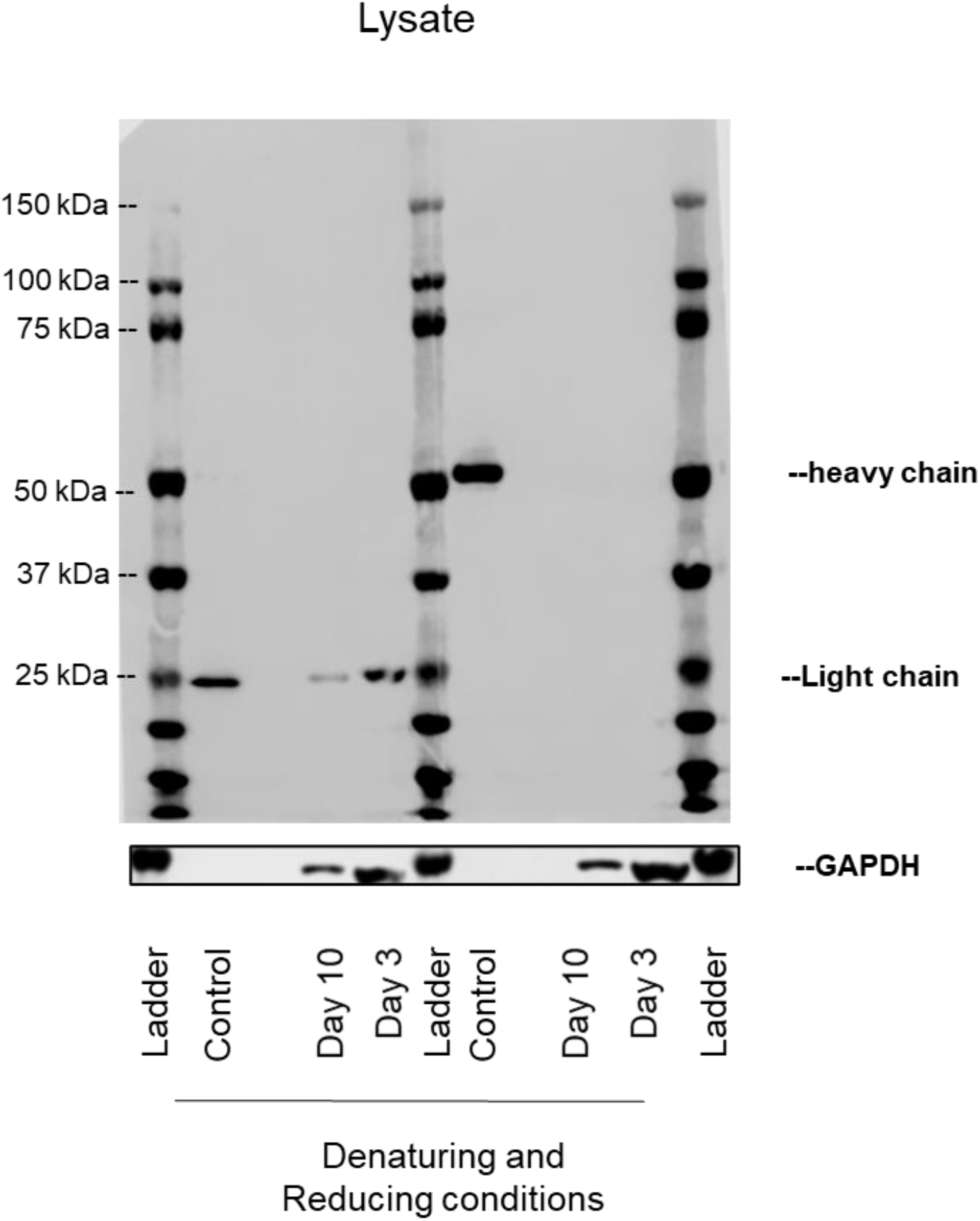
The light chain of the anti-IL8 antibody was detected in the lysate of the cells harvested after 3 and 10 days of culture. The samples were analyzed with Western blot under denaturing and reducing conditions. (full blot of figure 1b)

**Figure S3:**
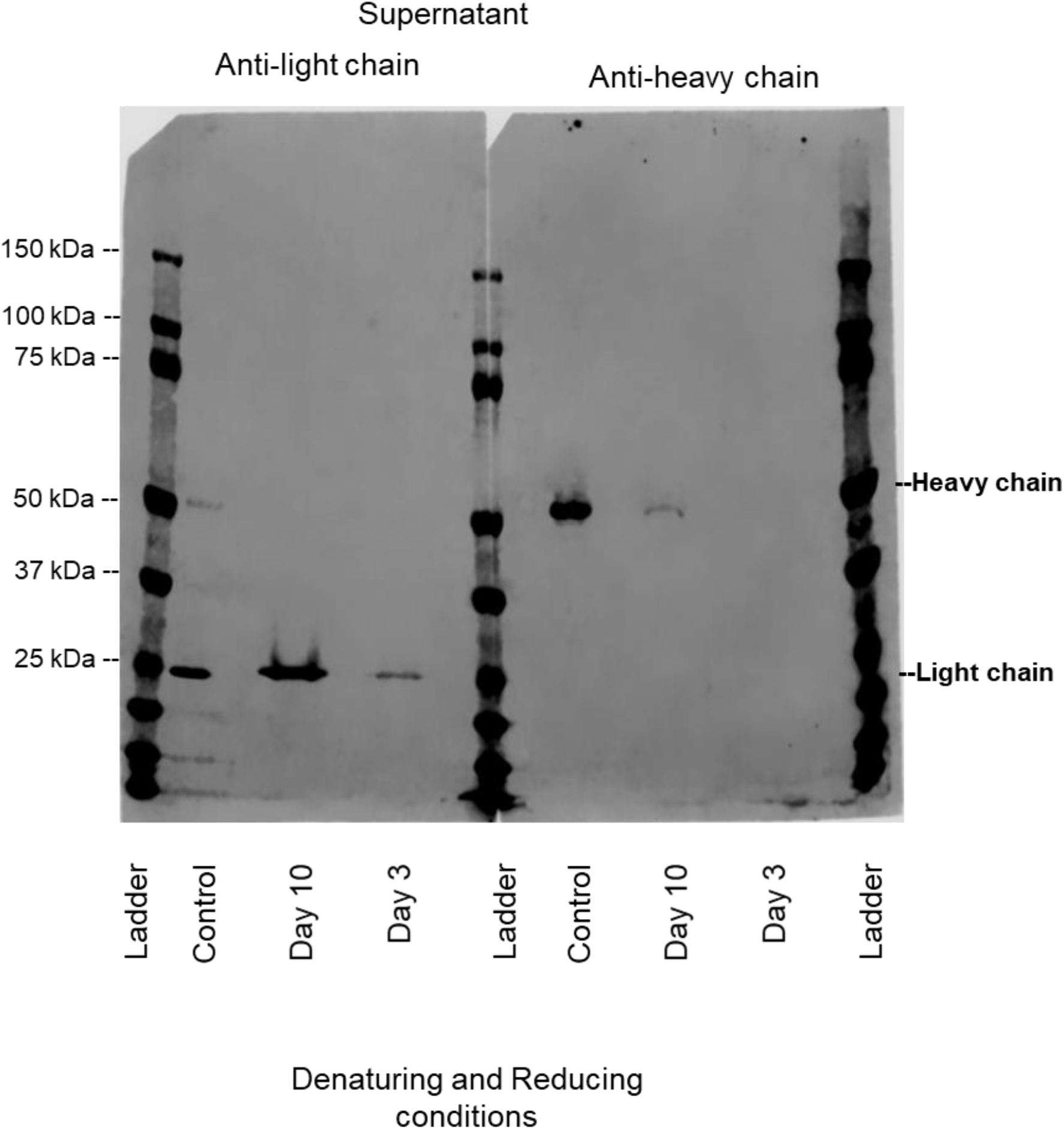
The heavy chain of the anti-IL8 antibody was detected in the supernatant only of the samples harvested after 3 and 10 days of culture. The samples were analyzed with Western blot under denaturing and reducing conditions. (full blot of figure 1c)

**Figure S4:**
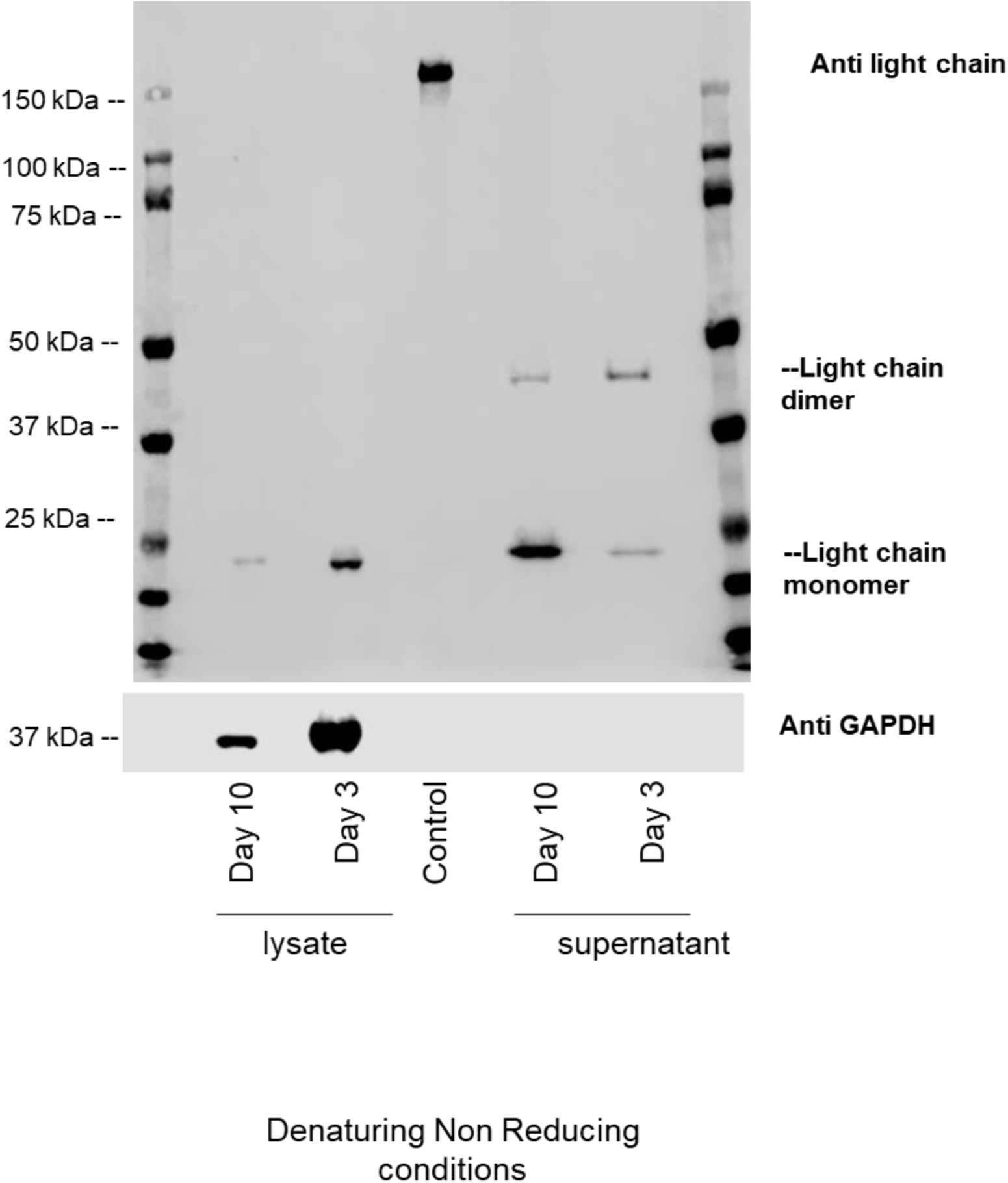
Western blot analysis of intracellular and extracellular light chains under denaturing non-reducing conditions revealed the presence of light chain dimers in the supernatant of cultures harvested after 3 and 10 days. (The figure shows the full blot of Figure 1d)

**Figure S5:**
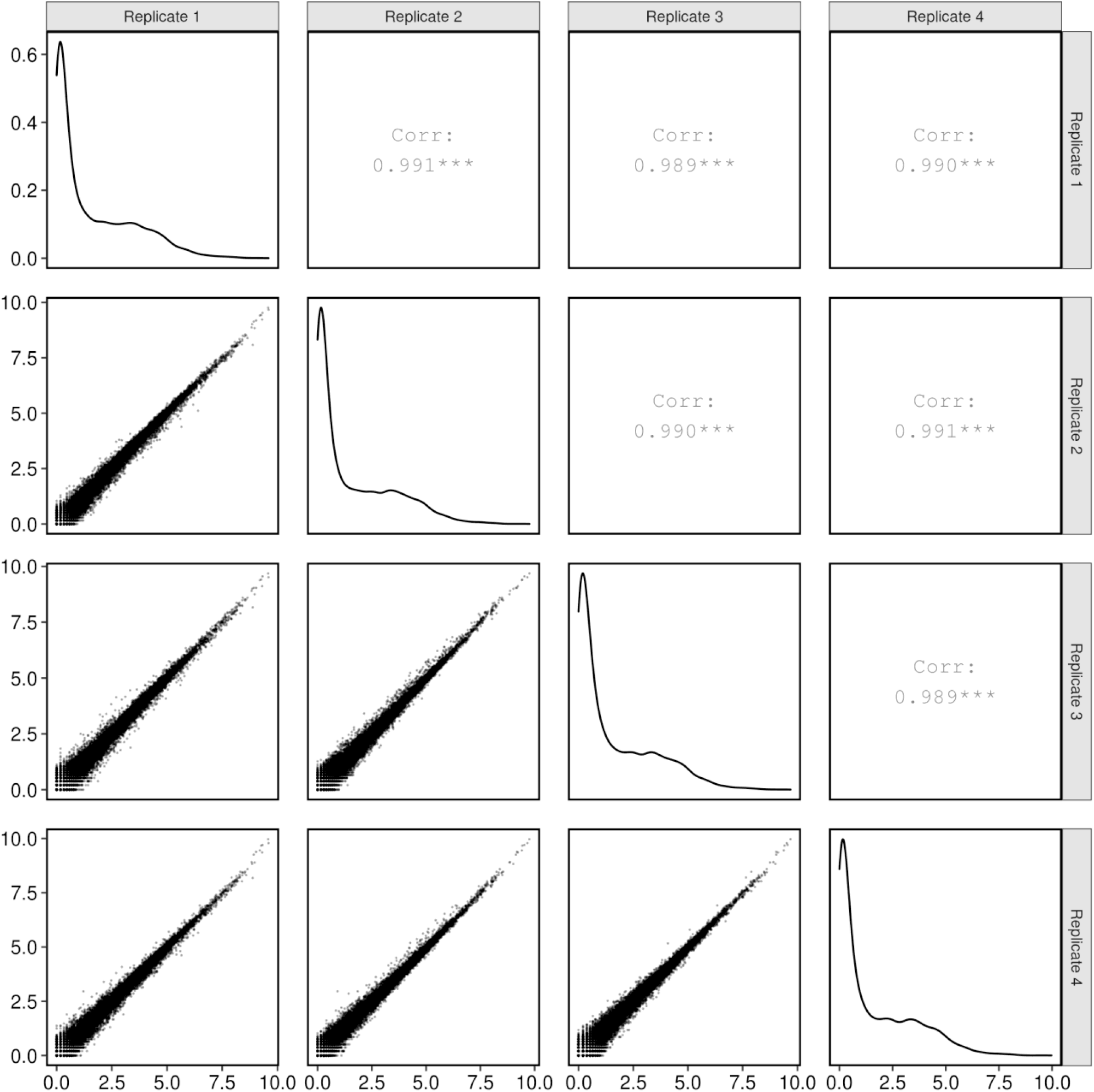
Correlation between CHO DP-12N1 scRNA-seq replicates. To determine the correlation between each of the replicate samples run on the BD Rhapsody WTA platform a “pseudobulk” sample was generated by summing the UMI counts for gene in each replicate and transforming to log2(TPM+1).

**Figure S6:**
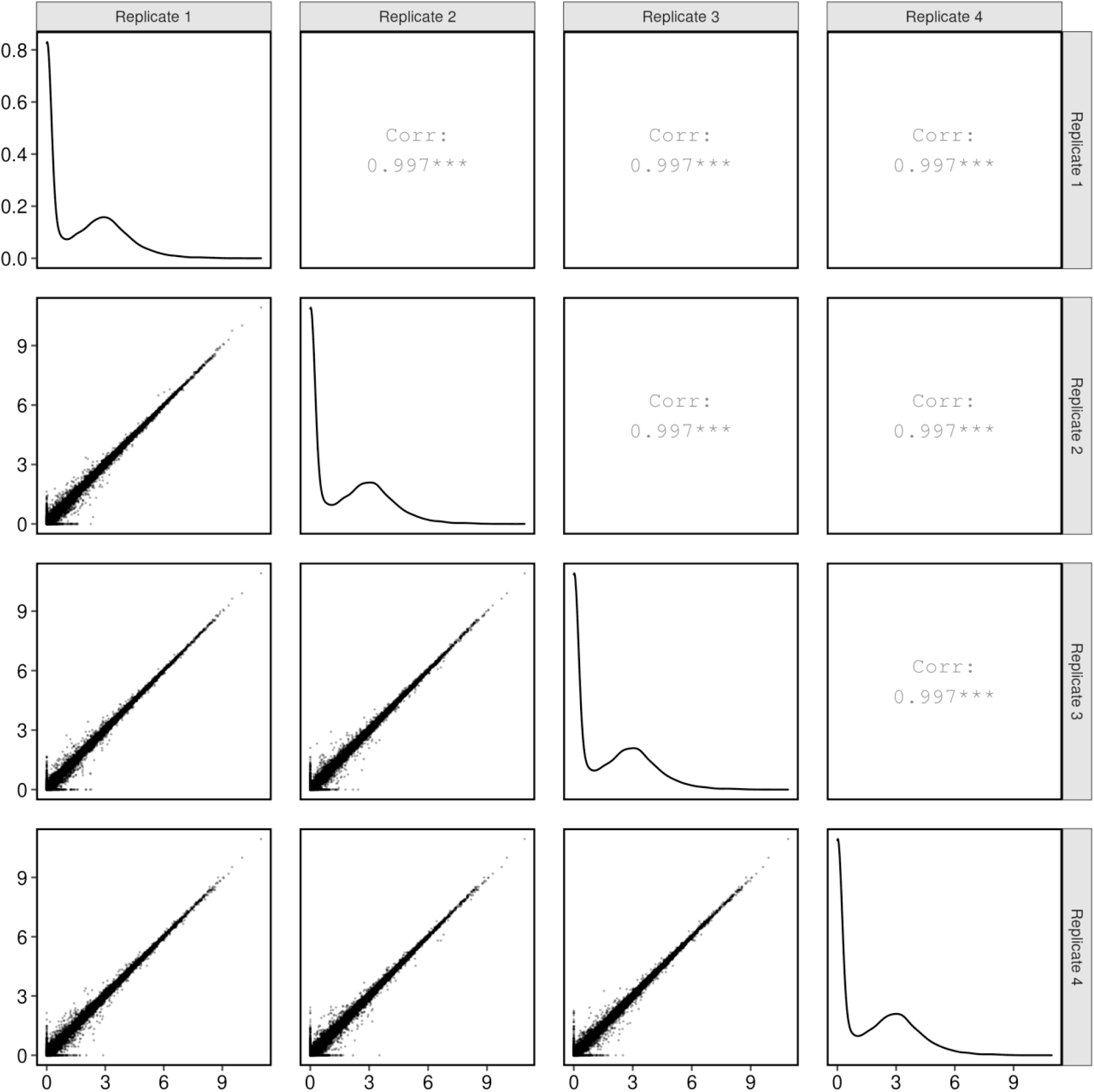
Correlation between CHO DP-12N1 bulk RNA-seq replicates. To determine the correlation between each of the replicate samples following bulk RNA-seq, Kalisto was used to calculated an aggregated TPM value for each gene. The plot shows the log2(TPM+1) for each sample.

**Figure S7:**
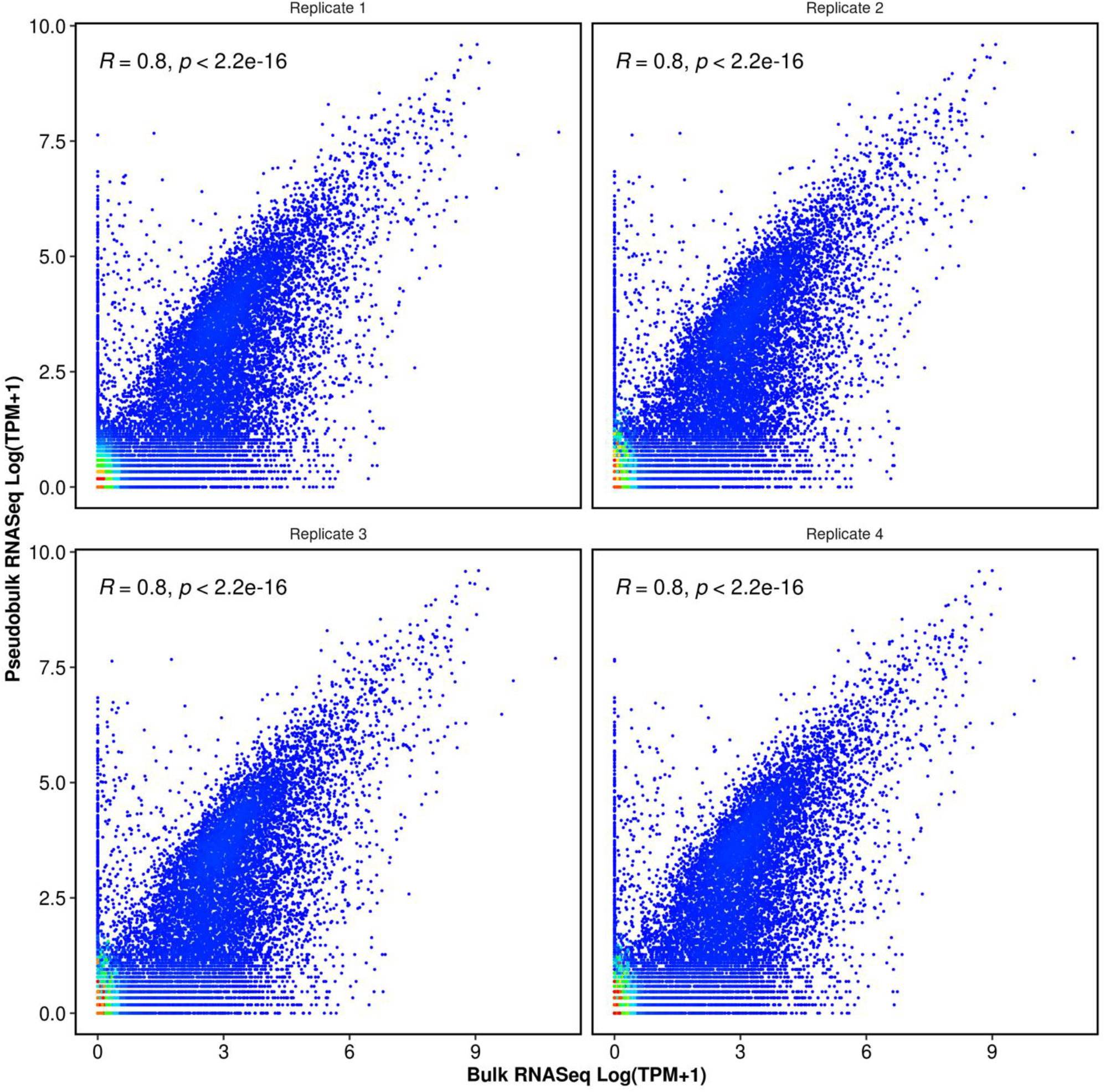
Correlation between scRNA-seq replicates on the Rhapsody system. To determine the agreement between each single cell RNA-seq replicate a pseudobulk expression sample was generated by summing the UMI counts for each gene before TPM scaling.

**Figure S8:**
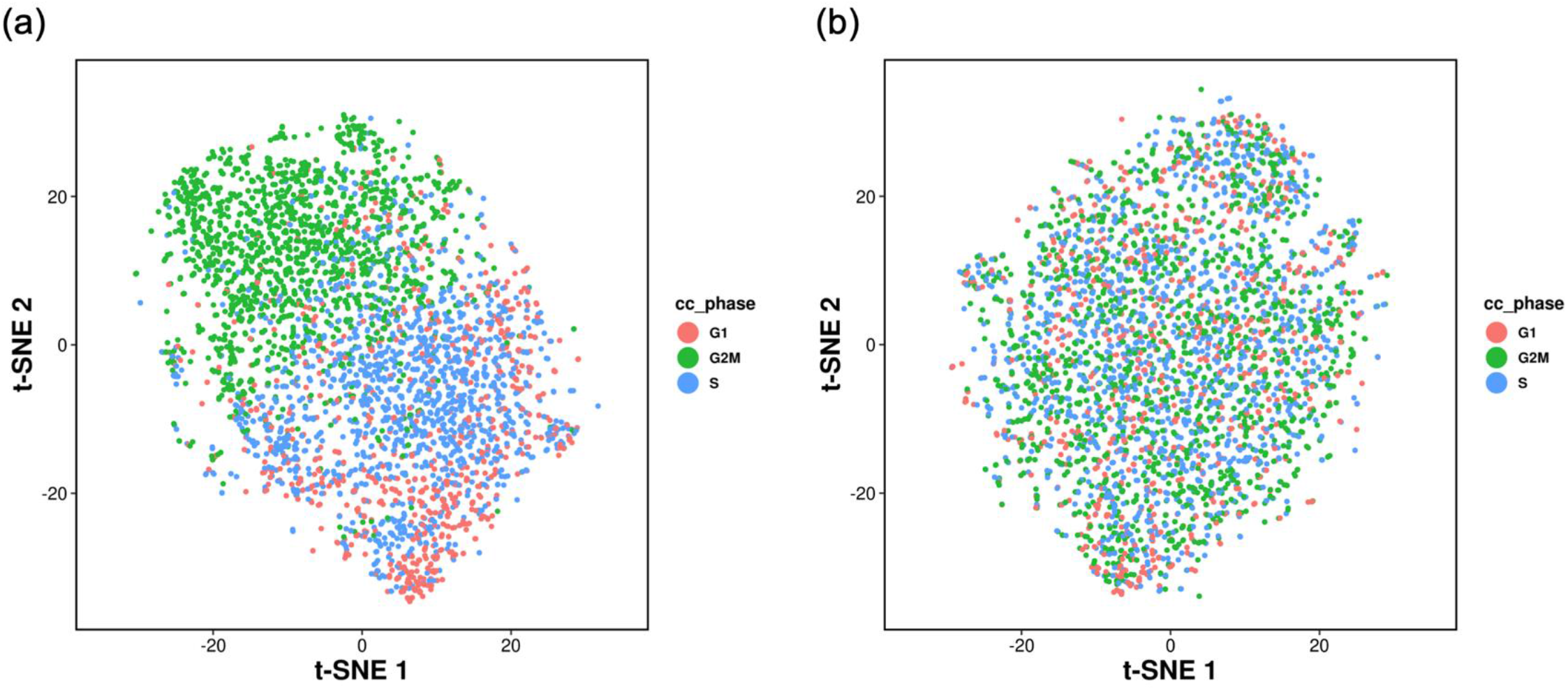
Correction of the effect of cell cycle in the CHO DP-12N1 scRNA-seq data. The Seurat Cell Cycle scoring method was used to assign a score indicating the probability that a cell was in either the S or G2/M phase of the cell based on the expression of known genes in each of these phases. To demonstrate the effectiveness of this approach principal components analysis was carried out using **(a)** uncorrected and **(b)** corrected data. The S and G2/M scores were used to ensure that the effect of cell cycle was reduced during t-SNE, trajectory and differential gene expression analyses conducted in Monocle.

**Figure S9:**
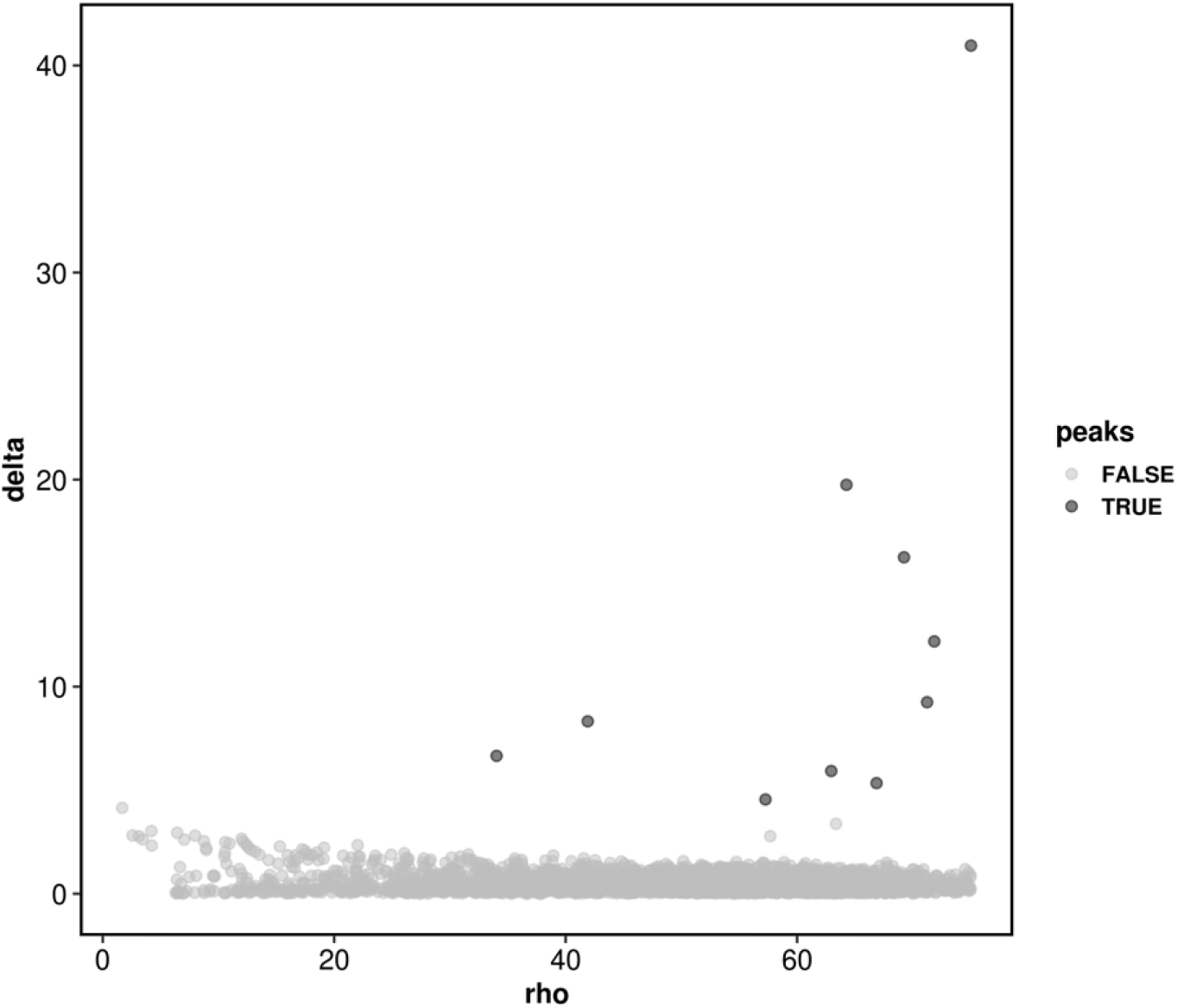
Selection of densityPeak clustering delta and Rho parameters.

**Figure S10:**
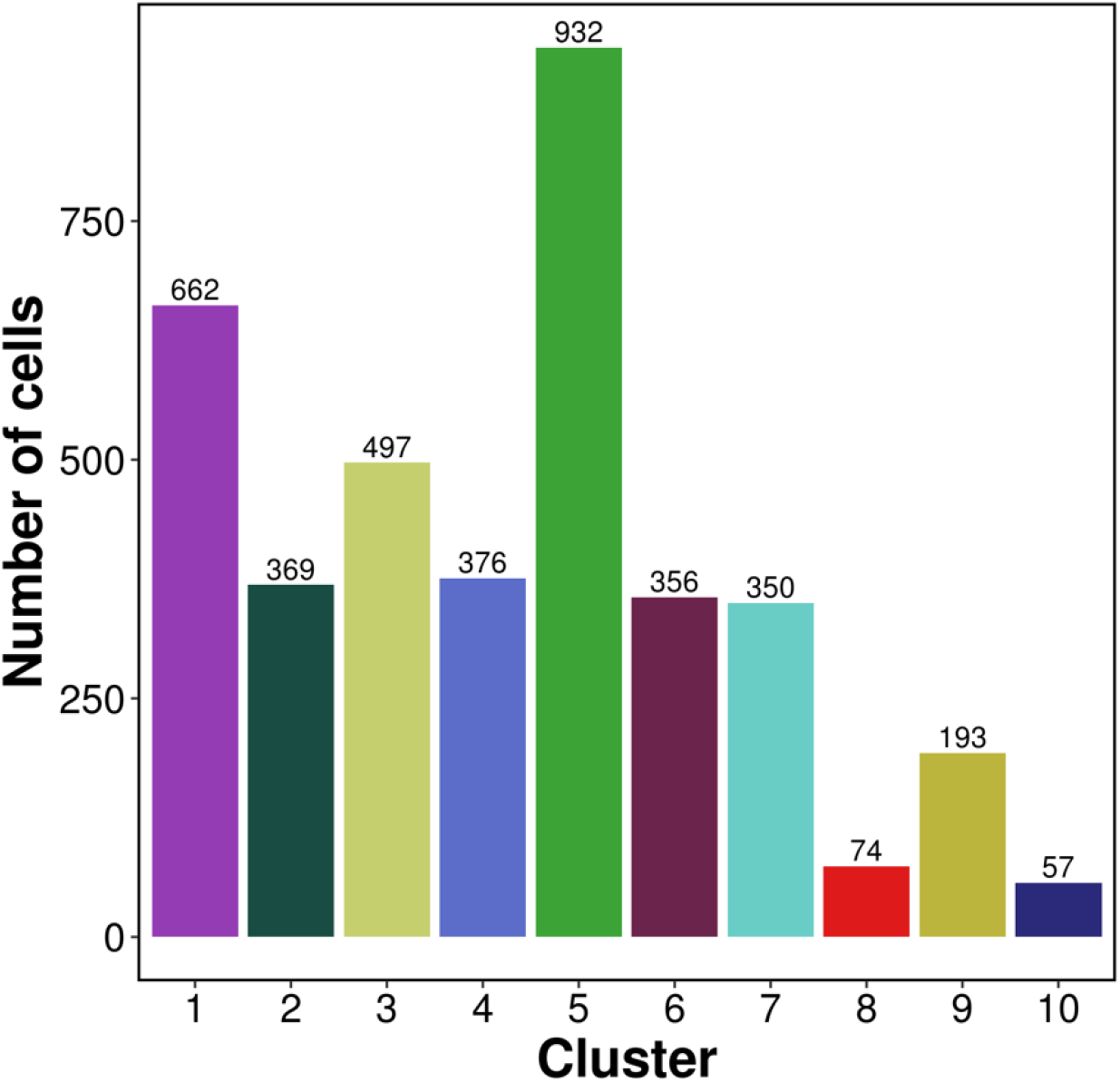
Cell numbers in each of the 10 clusters identified from t-SNE.

**Figure S11:**
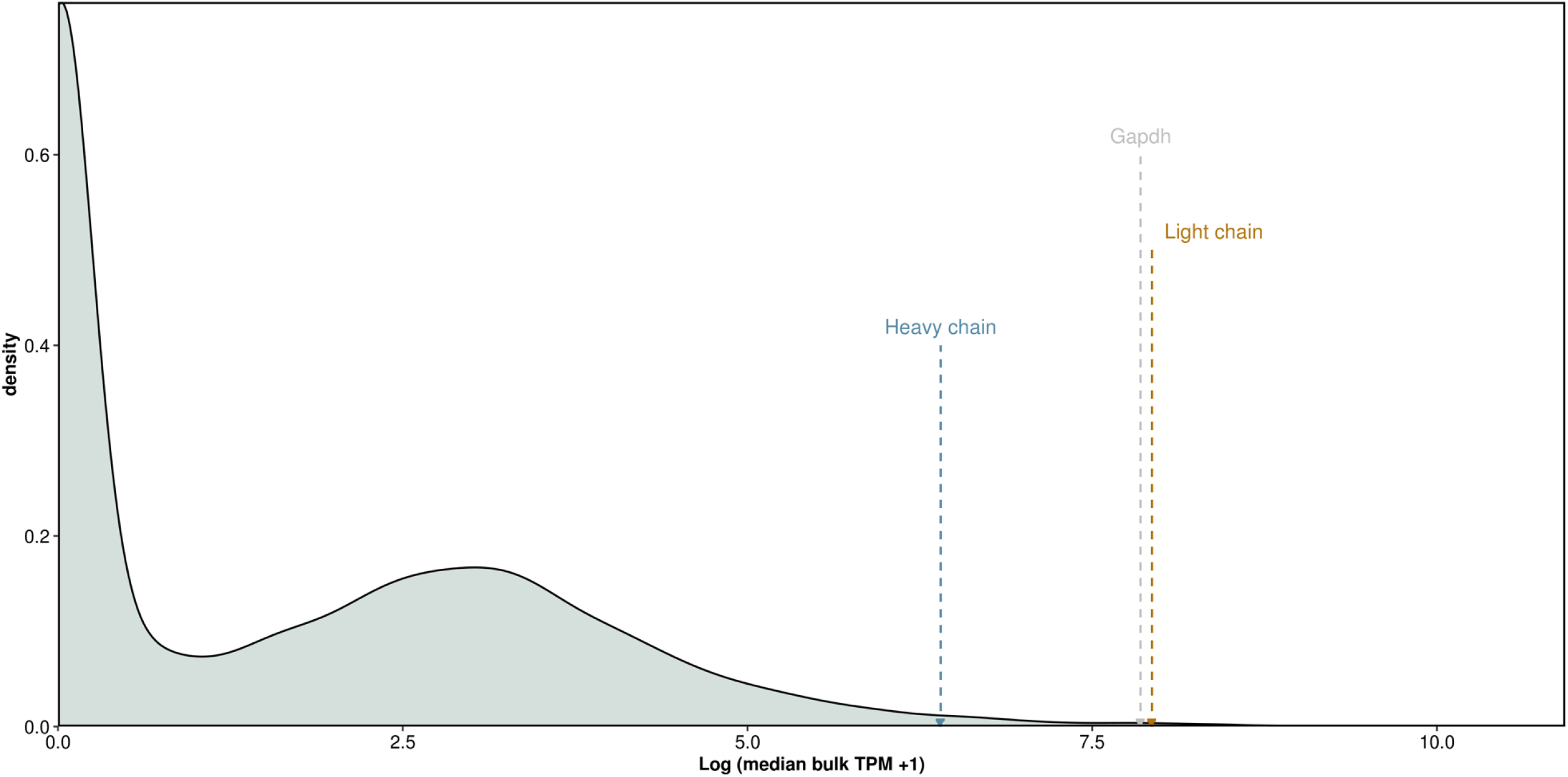
Bulk RNA-seq analysis of heavy and light chain gene expression. Similar to the single cell RNA-seq data the expression of the light chain was higher than that of the heavy chain. When compared to transcriptome both mAb genes where in the top 2% of all expressed genes.

## Notes

### Summary of Updates

Response to reviewers

